# Sucrose Non-fermenting Related Kinase (SNRK) Deficiency-Mediated Osteogenic Differentiation in Vascular Smooth Muscle Cells Exacerbates Vascular Calcification

**DOI:** 10.64898/2026.11.11.766061

**Authors:** Liu Ouyang, Chang-Jiang Yu, Wei-Dong Ye, Zhang Zheng-Zhipeng, Huan Nie, Cheng Jin-Kun, Xie Chen, Chang-Ming Xie, Zhi-Xue Liu, Xiao-Yan Su, Hui Huang, Ming-Hui Zou

**Author notes:** L.O., C.J.Y., and W.Y. contributed equally to this work. **Address for Correspondence:** Ming-Hui Zou, MD, PhD, 157 Decatur St SE, Atlanta, GA 30303, USA.; Hui Huang, MD, PhD, Shennan Middle Road, Shenzhen, 510275, China.

## Abstract

**Background:** The differentiation of vascular smooth muscle cells (VSMCs) into an osteogenic phenotype is a critical step in vascular calcification progression. Sucrose non-fermenting related kinase (SNRK), a member of the AMP-activated protein kinase (AMPK) family, has been implicated in diverse cellular functions and pathological conditions. However, whether SNRK is involved in VSMC osteogenic differentiation and calcification remains undefined.

**Methods:** Three mouse models were used to evaluate the contributions of SNRK in vascular calcification: Myh11-CreER^T2^-driven VSMC-specific SNRK knockout in C57BL/6J mice subjected to vitamin D3 overload or adenine diet, and *Apoe^-/-^* mice with VSMC (Myh11-CreER^T2^)-, endothelial cell (Cdh5-Cre)-, or macrophage (LysM-Cre)-specific SNRK knockout on a chronic western diet. The *in vitro* calcification was modeled by treating primary mouse aortic VSMCs and human aortic VSMCs (HASMCs) with phosphate (Pi).

**Results:** Cross-species transcriptomic analyses reveal reduced SNRK expression in murine vascular calcification and human calcified plaques. Human femoral artery samples from the elderly individuals with calcification exhibited reduced SNRK protein levels compared to non-calcified samples. In patients with chronic kidney disease, leukocyte levels of SNRK antisense RNA 1 (*SNRK-AS1*) were positively correlated with thoracic artery calcification Agatston scores. In parallel, decreased SNRK expression was observed in aortic arteries from three mouse models of calcification and in VSMCs treated with phosphate. Notably, VSMC-specific deletion of SNRK exacerbated calcific lesions, accompanied by upregulation of both runt-related transcription factor 2 (RUNX2) and msh homeobox 2 (MSX2). Conversely, SNRK deletion in either endothelial cells or macrophages had no impact on vascular calcification. Further, overexpression of SNRK in VSMCs, achieved via intravenous injection of an adeno-associated virus (AAV) ligated with VSMC *Tagln* promoter encoding *Snrk*, significantly attenuated vascular calcification along with reduced expression of osteogenic markers. Further interactome analysis demonstrated that SNRK directly binds to β-Catenin, a critical regulator of vascular calcification, through the amino acid sequence 270-765. Mechanistically, SNRK facilitated the interaction between β-Catenin and cullin 1/β-TrCP E3 ubiquitin ligase complex, promoting ubiquitin-proteasome-mediated degradation of β-Catenin and subsequently reducing the expression of osteogenic markers in VSMCs.

**Conclusions:** We conclude that SNRK safeguards against vascular calcification by preventing β- Catenin/RUNX2/MSX2-mediated VSMC osteogenic differentiation.

**Clinical Perspective:** *What is new?:*

- This study reveals the previously unrecognized protective role of sucrose non-fermenting related kinase (SNRK) in vascular calcification, vascular smooth muscle cell defective of SNRK result in enhanced β-Catenin, runt-related transcription factor 2 (RUNX2), and msh homeobox 2 (MSX2) and ectopic calcium deposition within vessel wall.
- SNRK exerts potent inhibitory effects on osteogenic activity in vasculature by promoting E3 ligase complex formation between β-TrCP and cullin 1, thus driving ubiquitin-proteasome-mediated degradation of β-Catenin.
- Elderly individuals with femoral artery calcification have impaired SNRK levels in vascular smooth muscle cell layer. Chronic kidney disease patients with thoracic artery calcification have increased leukocyte SNRK antisense RNA 1 (*SNRK-AS1*) level, and *SNRK-AS1* level is associated with calcification development in the clinical setting.

*What are the clinical implications?:*

- Targeting SNRK/β-Catenin/RUNX2/MSX2 axis represents a promising therapeutic strategy for treating vascular calcification.
- The identification of *SNRK-AS1* as a potential biomarker provides a valuable approach for diagnosis and risk stratification in populations vulnerable to vascular calcification.

## Introduction

Vascular calcification, characterized by the pathological deposition of calcium nodules in blood vessels, is common in chronic kidney disease, atherosclerosis, type II diabetes, and aging, and it significantly increases the risk of myocardial infarction and stroke.^1-5^ Current therapies target mineral imbalances using pyrophosphate analogs, fetuin-A, and vitamin K-dependent proteins, or employ invasive procedures like atherectomy.^6-9^ However, these approaches are limited in preventing early calcium deposition.

Vascular smooth muscle cells (VSMCs), the primary cells in the arterial medial layer, play a pivotal role in vascular calcification through an active process resembling bone formation.^10,11^ Under stimulus such as disrupted phosphate homeostasis,^12^ inflammatory factors,^13^ oxidative DNA damage,^14^ and matrix remodeling,^15^ VSMCs undergo osteogenic differentiation. This shift, marked by the loss of VSMCs contractile markers and activation of osteogenic factors like runt-related transcription factor 2 (RUNX2), msh homeobox 2 (MSX2), and β-Catenin, promotes the deposition of calcium-phosphate crystals in the extracellular matrix, contributing to disease progression.^16^ However, the endogenous mechanisms suppressing VSMCs osteogenic differentiation and calcification remains poorly defined.

Sucrose non-fermenting-related kinase (SNRK) belongs to the AMP-activated protein kinase (AMPK)-related superfamily of serine/threonine kinases and can be phosphorylated by liver kinase B1 (LKB1) at the conserved T-loop threonine residue (T173).^17-20^ SNRK participates in various pathological conditions, including neuronal apoptosis,^21^ tumorigenesis,^22,23^ and inflammatory response.^24-27^ Notably, SNRK has been implicated in cardiovascular physiology and pathology. In mice, global SNRK knockout causes neonatal lethality with cardiac enlargement, conditional deletion in cardiomyocytes leads to heart failure.^28^ SNRK regulates cardiac metabolic homeostasis through phosphorylation of Rho-associated kinase and mitigates ischemia-reperfusion injury by inhibiting UCP3-mediated mitochondrial dysfunction.^29,30^ In addition, SNRK reduces DNA damage responses, protecting against cardiac hypertrophy.^31^ Furthermore, studies have shown that SNRK is essential for arterial specification in zebrafish, and mediating hypoxia-induced angiogenesis in mice.^32-34^ These findings underscore the pivotal role of SNRK in maintaining cardiovascular homeostasis. However, whether SNRK contributes to VSMC pathophysiology, particularly vascular calcification, remains unknown.

In this study, we demonstrated that the deletion of SNRK in VSMCs, but not in endothelial cells or macrophages, significantly exacerbates VSMC osteogenic differentiation and subsequent calcification. Our results suggest that SNRK may serve as a promising therapeutic target for preventing and treating vascular calcification.

## Methods

The data supporting the conclusions of this study can be obtained from the corresponding author upon reasonable request. Detailed methods are available in the supplemental material.

### Study population

This case-sectional study was conducted at Tungwah Hospital, China, between March 2018 and January 2020. A total of 198 participants with chronic kidney disease (CKD) were initially enrolled as primary cohort. However, 10 participants were excluded from meeting the exclusion criteria, another 10 for missing peripheral blood collection, and 16 for lacking thoracic artery calcification data obtained through multidetector computed tomography (MDCT) scans. Written informed consent was obtained from all participants prior to their enrollment. For external validation, using the same inclusion and exclusion criteria, we collected an independent clinical cohort from another center between November 2023 and December 2025, the Eighth Affiliated Hospital of Sun Yat-sen University, consisting of a total of 110 patients with chronic kidney disease (CKD) from the Department of Nephrology. The study adhered to the ethical guidelines of the 1975 Declaration of Helsinki and received approval from the Ethics Committee of the Tungwah Hospital (2017DHLL017) and the Eighth Affiliated Hospital of Sun Yat-sen University (2023-100-01).

### Animal experiments

All animal experiments were approved by the Georgia State University Institutional Animal Care and Use Committee and conducted in accordance with the NIH Guide for the Care and Use of Laboratory Animals. Specific-pathogen-free mice were housed in the animal facility of Georgia State University under controlled conditions, including uniform temperature, humidity, and a 12-hour light-dark cycle. The mice had unrestricted access to sterilized water and the designated diet. Mice were anesthetized with isoflurane (5% for induction, inhalation) until loss of reflex, followed by cervical dislocation for euthanasia and subsequent tissue collection.

### Statistical analyses

All experimental results were expressed as mean ± standard error of the mean (SEM), and clinical results as mean ± standard deviation (SD), median (25th-75th quartiles) for continuous variables, or n (%) for categorical variables, respectively. The Shapiro-Wilk test or D’Agostino-Pearson test was performed to assess data normality. The equality of group variances was assessed using *F* test or Brown-Forsythe test. For comparisons between two groups, significance was assessed using unpaired two-tailed Student’s *t*-tests, unpaired *t*-test with Welch’s correction, or nonparametric Mann-Whitney *U* test. For comparisons among multiple groups, analysis of variance (ANOVA) or Welch’s ANOVA test accompanied by Tukey’s post hoc test (equal variances assumed), Dunnett’s post hoc test, or Sidak post hoc test (equal variances not assumed) was performed.

Correlation was assessed using Spearman’s rank correlation coefficient. Model discrimination was assessed using the c-Statistic (AUC), with differences between models evaluated by the DeLong test. Reclassification was assessed using integrated discrimination improvement (IDI) and continuous net reclassification improvement (NRI). Missing values were imputed using multiple imputations. Analyses were subsequently conducted on the imputed datasets. All statistical analyses were performed by two independent researchers in a blinded manner. *P* < 0.05 was considered statistically significant. Statistical analyses were performed using SPSS version 29.0 (IL, USA) or GraphPad Prism 9.5 (CA, USA).

## Results

### Reduced SNRK expression in murine vascular calcification and human calcified plaques revealed by bulk and single-cell transcriptomics

We screened bulk RNA-seq samples of healthy and calcified mouse aorta from the GEO database. Compared with healthy non-calcified aorta samples, we noticed that *Snrk* expression was significantly decreased in calcified aorta (Figure 1A and Figure S1). To further investigate the correlation between SNRK and calcification in human, we analyzed single-cell transcriptome data from calcified atherosclerotic core (AC) plaques and proximal adjacent (PA) portions. After removing low-quality data, a total of 50,538 cells were obtained, including AC and PA cells from three patients. Through batch effect correction, dimensionality reduction, and clustering, all cells were divided into 17 cell clusters (Figure 1B and Figure S2). Based on previously reported marker genes in the literature, these cell clusters were annotated into different cell types, and the proportions of VSMCs were reduced in AC (Figure 1C-1E). We found that among the four VSMC subclusters, SNRK exhibited differential expression only in the VSMC-1 subcluster (Figure 1F). In VSMC-1, the expression level of *SNRK* in AC was significantly lower than that in PA (*P* = 1.7e-21).

**Figure 1.**
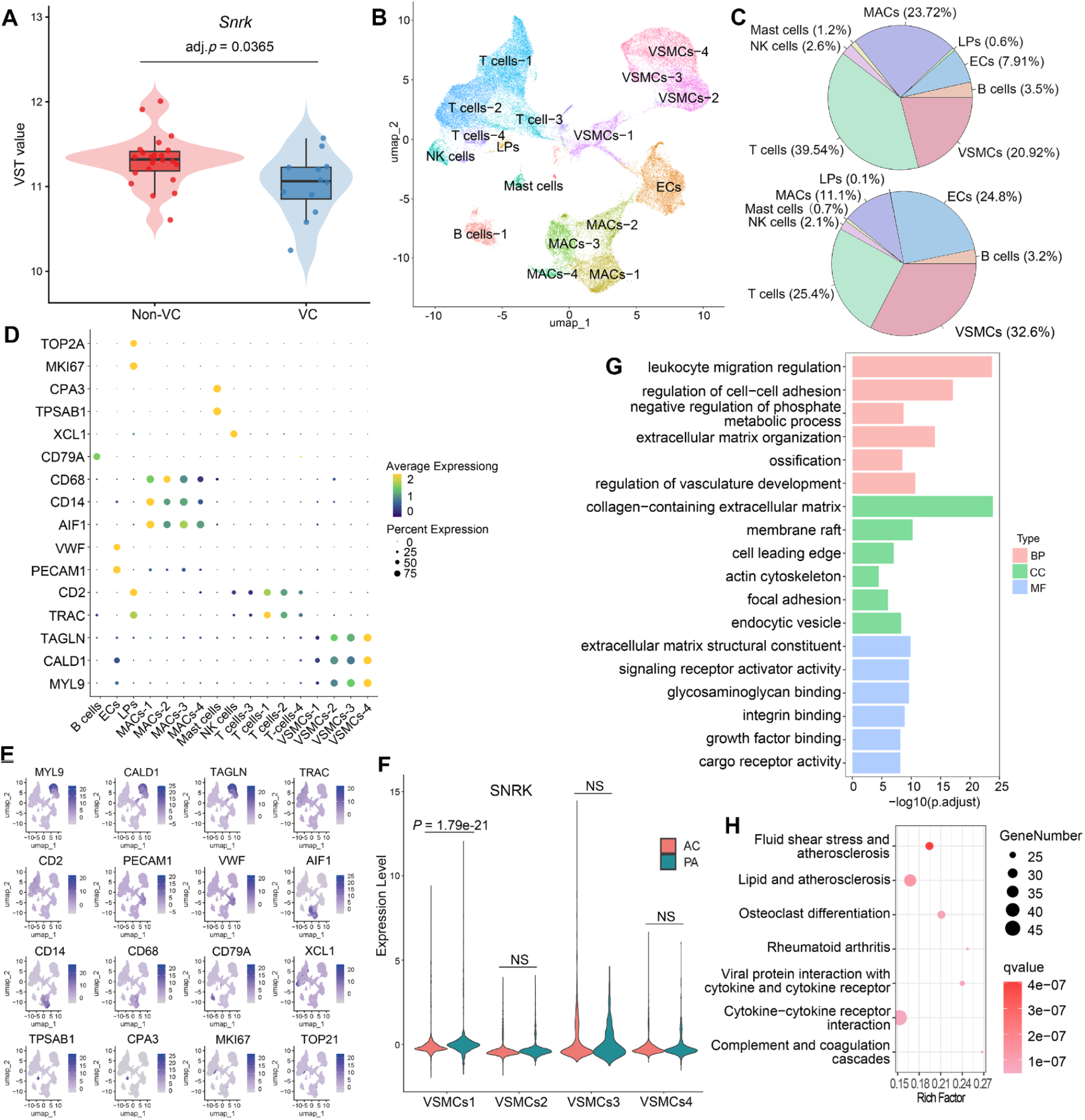
Cross-species transcriptomic analyses reveal reduced SNRK expression in vascular calcification. (**A**) Violin plot showing variance-stabilized (VST) expression of *Snrk* in murine aortic samples from non-vascular calcification (Non-VC) and vascular calcification (VC) groups. (**B**) UMAP visualization of 17 cell types in calcified atherosclerotic core (AC) and patient-matched proximal adjacent (PA) cells after batch effect correction. (**C**) Proportions of different cell types in AC (upper panel) and PA (lower panel). (**D**) Bubble plot displaying marker genes of each cell cluster. The size of the dots represents the percentage of cells expressing the marker gene in the cluster, and the yellower the dot color, the higher the expression level. (**E**) UMAP visualization of the expression level distribution of marker genes in cells; the darker the color, the higher the expression level. (**F**) Expression differences of *SNRK* in the four VSMC subclusters between AC and PA. NS indicates no significant difference. (**G**) Bar plot of GO enrichment results for DEGs in VSMC-1 between AC and PA. (**H**) Bubble plot of KEGG enrichment results for DEGs in VSMC-1 between AC and PA; the darker the bubble color, the higher the enrichment level.

We performed functional enrichment analysis of Differentially Expressed Genes (DEGs) in VSMC-1 between AC and PA. The results showed that these DEGs were significantly enriched in a series of terms and pathways related to calcific phenotype transition of VSMC, such as fluid shear stress and atherosclerosis, lipid and atherosclerosis, ossification, and osteoclast differentiation (Figure 1G and 1H). Taken together, cross-species transcriptomic analyses suggest that SNRK are involved in regulating the development of vascular calcification.

### SNRK reduction in humans and three mouse models of calcification correlates with both a loss of contractile VSMC marker and enhanced osteogenic factors

To explore the clinical relevance of SNRK expression and vascular calcification, we analyzed autopsy samples from elderly individuals, performing histological and immunofluorescence staining in the same area. The results revealed that SNRK levels, along with the contractile VSMC marker α-SMA, were significantly decreased in calcified lesion compared to those without calcification, as confirmed by von Kossa staining (Figure 2A, Figure S3).

**Figure 2.**
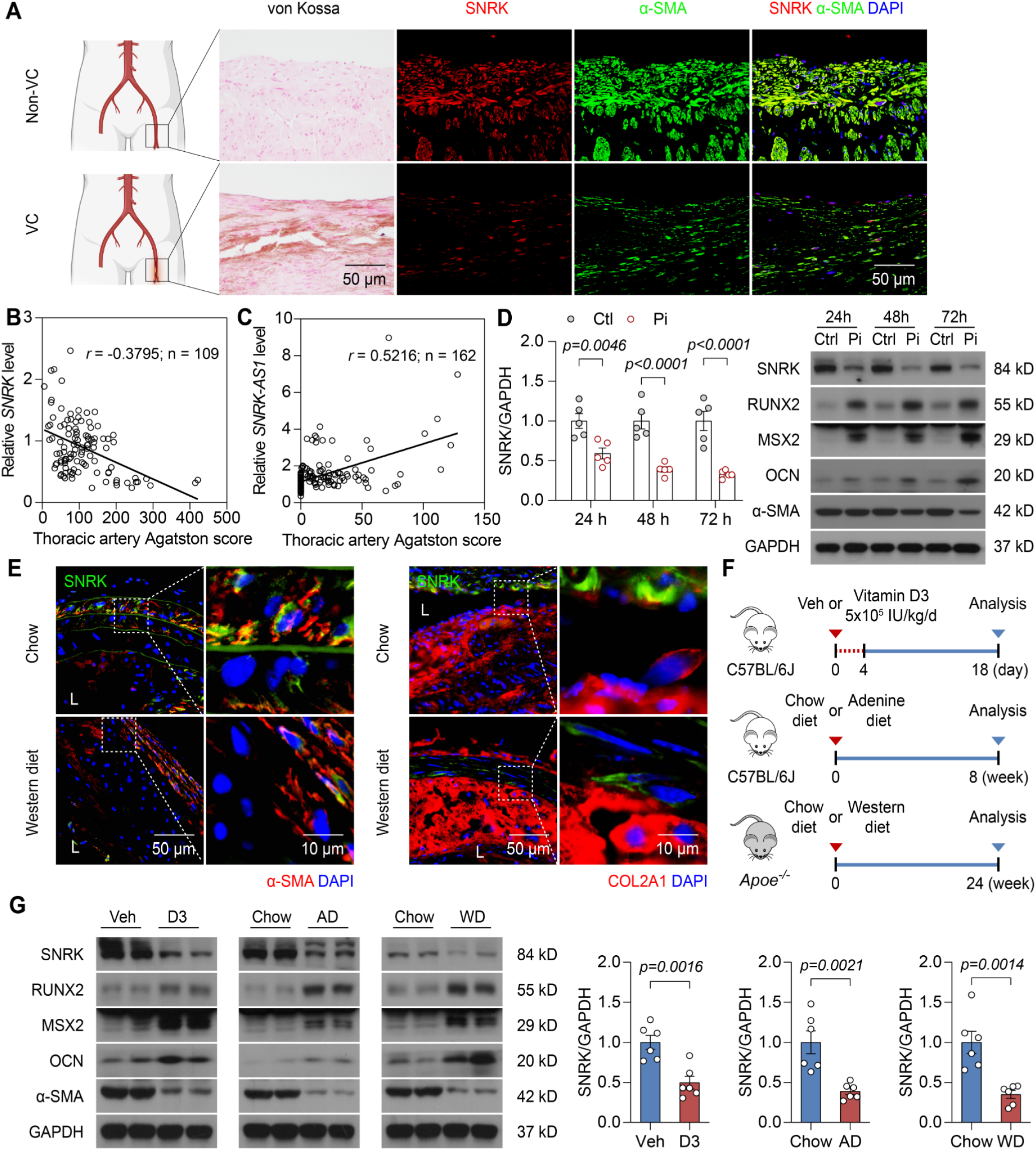
SNRK reduction in humans and three mouse models of calcification correlates with both a loss of contractile VSMC marker and enhanced osteogenic factors. (**A**) Representative von Kossa staining and immunofluorescence staining of SNRK in sequential section of non-calcified (Non-VC) or calcified (VC) human femoral artery (n = 4-6). Scale bar: 50 μm. (**B**) The correlation of PBMC *SNRK* level and square rooted thoracic artery Agatston score in CKD patients of external validation cohort. (**C**) The correlation of PBMC *SNRK-AS1* level and square rooted thoracic artery Agatston score in CKD patients of primary cohort. (**D**) Western blot analysis of SNRK and VSMC markers in human VSMCs (HASMCs) treated phosphate (Pi, 2.6 mmol/L) for indicated time (n = 5). (**E**) Immunofluorescence staining of SNRK co-located with α-SMA or COL2A1 in brachiocephalic artery of *Apoe*^-/-^ mice fed with chow diet or western diet for 24 weeks (n = 7-8). Lumen (L). Scale bars: 50 μm (left panels) and 10 μm (right panels). (**F**) Experimental scheme of three mouse models of vascular calcification. (**G**) Western blot analysis of SNRK and VSMC markers in aortic SMC layer of indicated mice (n = 6). Vitamin D3 (D3), Adenine diet (AD), Western diet (WD). Results are presented as mean ± SEM, and analyzed using Spearman’s rank correlation coefficient test for **B** and **C**, two-tailed unpaired Student’s *t*-test for **D** and **G**.

SNRK antisense RNA 1 (*SNRK-AS1*), a long non-coding RNA transcribed from the opposite strand of the *SNRK* gene, was identified as a regulator of SNRK expression. In the *SNRK-AS1* deletion human aortic VSMC (HASMC), both SNRK mRNA and protein expression levels were elevated, confirming that SNRK is targeted by *SNRK-AS1* (Figure S4). Analysis of peripheral blood mononuclear cells (PBMCs) mRNA from CKD patients revealed that *SNRK* expression inversely correlated with thoracic artery calcification (*r* = -0.3795, 95% CI -0.550 to -0.169; Figure 2B), whereas elevated *SNRK-AS1* level positively correlated with calcification (*r* = 0.5216, 95% CI 0.393-0.636; Figure 2C). Multiple linear regression analysis, adjusted for demographic and clinical covariates, indicated a significant association between PBMC *SNRK-AS1* levels and thoracic artery calcification in CKD patients (Table S1 and S2). Adding SNRK-AS1 to the conventional model significantly improved discrimination and reclassification performance, as evidenced by an increase in AUC from 0.812 to 0.927 ( ΔAUC 0.115; 95% CI 0.061-0.169; *p* < 0.001), an integrated discrimination improvement (IDI) of 0.258 (95% CI 0.190–0.327, *p* < 0.001), and a continuous Net Reclassification Improvement (NRI) of 1.21 (Table S3). These findings were consistently replicated in an independent external validation cohort, in which PBMC *SNRK* and *SNRK-AS1* remained independently associated with calcification (Figure S5, Table S4-S6). These findings suggest that SNRK and its regulation by *SNRK-AS1* correlate with the progression of vascular calcification.

VSMC calcification, similar to skeletal mineralization, is a complex process driven by several intracellular pathways, which are regulated by key transcription factors like RUNX2 and MSX2. To determine whether these factors were involved in our model, we induced VSMC calcification by exposing cells to osteogenic medium containing high phosphate (Pi, 2.6 mmol/L). After 24 hours, osteogenic markers RUNX2, MSX2, and osteocalcin (OCN) were significantly upregulated in comparison to control HASMCs (Figure 2D, Figure S6). In contrast, both SNRK protein level and α-SMA expression were notably reduced in the calcified VSMCs (Figure 2D).

To further investigate the relationship between SNRK and vascular calcification *in vivo*, we examined brachiocephalic arteries from *Apoe^-/-^*mice fed either a chow or a western diet for 24 weeks. Immunofluorescence analysis revealed decreased SNRK protein levels in the calcified aortas of the *Apoe^-/-^*mice, as evidenced by an increase in osteogenic marker Collagen II alpha1 (COL2A1) expression and a reduction in α-SMA and (Figure 2E, Figure S7).

Additionally, we generated a chronic renal failure mouse model by feeding the mice an adenine-rich diet for 8 weeks to induce vascular calcification as previously described.^35^ We also administered 5x10^5^ IU of vitamin D3 per kilogram intraperitoneally for 4 consecutive days to further induce vascular calcification (Figure 2F). Both models exhibited elevated calcium deposition in the aortas, confirming successful vascular calcification induction (Figure S8). In these calcified aortas (with the endothelium and adventitia removed), SNRK and α-SMA levels were reduced, while RUNX2, MSX2, and OCN levels were increased (Figure 2G, Figure S9). Collectively, these findings suggest that reduced SNRK expression may be linked to vascular calcification.

### SNRK regulates VSMC osteogenic reprogramming

Osteogenic reprogramming of VSMC is the central event in vascular calcification. Effects of SNRK on VSMC osteogenic reprogramming and calcification were further verified *in vitro* and *in vivo*. The mRNA levels of osteogenic markers including *RUNX2*, *MSX2*, *SOX9*, *ALPL*, *SPP1*, *and COL1A1* exhibited significant increase in SNRK-silenced HASMCs (Figure 3A). Alizarin red staining for calcified nodules indicated calcification in HASMCs stimulated by high-Pi with SNRK deficiency (Figure 3B). The effect of SNRK on calcification was further verified in HASMCs transfected with lentivirus carrying scramble shRNA or *SNRK* shRNA by osteogenic protein levels (Figure 3C). SNRK-loss-of-function HASMC underwent osteogenic reprogramming and became more susceptible to calcifying stimuli. *Snrk^Flox^* mice were crossed with *Myh11-CreER^T2^*mice to generate tamoxifen-inducible VSMC-specific SNRK knockout (*Snrk^△ SMC^*) mice. *Snrk^△ SMC^* mice were further crossed with hyperlipidemic apolipoprotein E-deficient (*Apoe^-/-^*) mice to generate *Apoe^-/-^ Snrk^△SMC^* (*Apoe^-/-^*-*Snrk^flox/flox^-Myh11-CreER^T2+^*) mice and *Apoe^-/-^ Snrk^Flox^* (*Apoe^-/-^*-*Snrk^flox/flox^-Myh11-CreER^T2-^*) mice. We observe SNRK deficiency amplified adenine diet-induced VSMC osteogenic reprogramming in calcified aortas, as evidenced by elevated MSX2, RUNX2, and OCN expression levels, in parallel with decreased α-SMA protein level (Figure 3D). Consistently, immunofluorescence staining revealed markedly increased RUNX2, MSX2, and COL2A1 in brachiocephalic artery of western diet-induced *Apoe^-/-^ Snrk^△SMC^* mice, compared with their littermates (Figure 3E).

**Figure 3.**
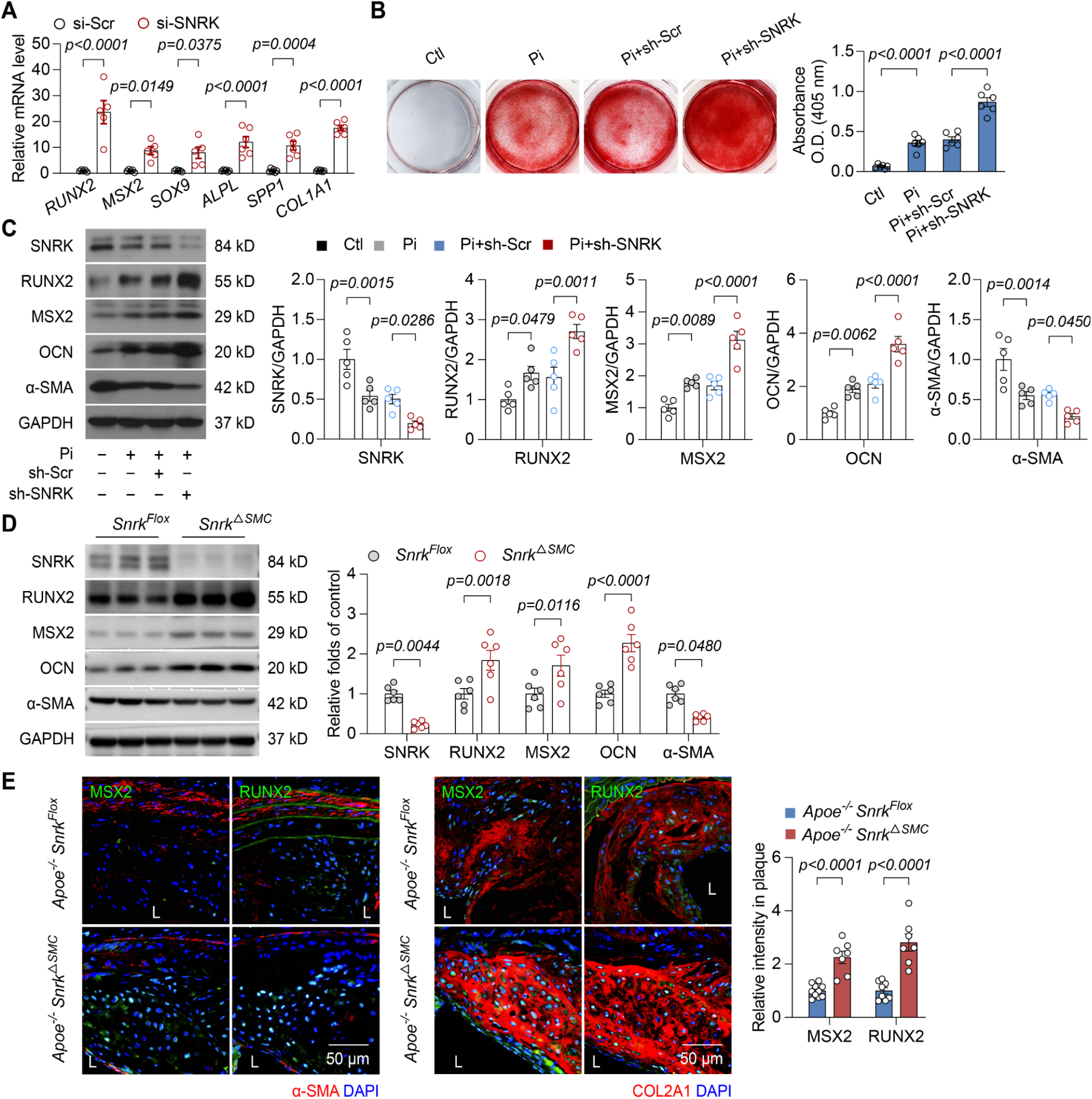
SNRK regulates VSMC osteogenic reprogramming. (**A**) RT-PCR analysis of osteogenic reprogramming genes in phosphate (Pi, 2.6 mmol/L)-treated HASMCs transfected lentivirus of scramble (si-Scr) or SNRK siRNA for 24 hours (n = 5-6). (**B**) Representative whole-well alizarin red S staining of Pi-treated HASMCs infected lentivirus of scramble (sh-Scr) or *SNRK* shRNA for 14 days. Calcium deposition was quantified by measuring absorbance at 405 nm after dye extraction. Data are presented as relative calcium content (n = 6). (**C**) Western blot analysis of SNRK and VSMC marker genes in Pi-treated HASMCs transfected lentivirus of scramble (sh-Scr) or *SNRK* shRNA for 14 days (n = 5). (**D**) Western blot analysis of SNRK and VSMC markers level in aortas of *Snrk* Flox (*Snrk^Flox^*) or VSMC-specific *Snrk* knockout mice (*Snrk^△SMC^*) mice fed with adenine diet for 8 weeks (n = 6). (**E**) Immunofluorescence staining of MSX2 and RUNX2 co-located with α-SMA or COL2A1 in brachiocephalic artery of *Apoe*^-/-^ *Snrk^Flox^* or *Apoe*^-/-^ *Snrk^△SMC^* mice fed with western diet for 24 weeks (n = 7-10). Lumen (L). Scale bars: 50 μm. Results are presented as mean ± SEM. *P* values are assessed using two-tailed unpaired Student’s *t*-test for **A** and **D**, one-way ANOVA with Sidak’s post hoc test for **B** and **C**.

### SNRK deficiency aggravates vascular calcification *in vivo*

To further assess the effect of SNRK on vascular calcification *in vivo*, we established three mouse models of calcification using VSMC-specific SNRK knockout mice and their littermates. Aortas isolated from *Snrk^△SMC^* mice treated with either vitamin D3 or adenine diet, and *Apoe*^-/-^ *Snrk^△SMC^* mice fed with western diet exhibited increased calcium content, compared with their littermates (Figure 4A and 4B). To ensure consistency, staining was performed on consecutive sections obtained from the same anatomical level of the vessel. More severe vascular calcification was confirmed by the increased intensity of von Kossa staining in the thoracic aorta, femoral artery, aortic arch, carotid artery, abdominal aorta, and iliac artery of *Snrk^△SMC^* mice. Notably, this was accompanied by dilated aortic diameter, due to the loss of vascular elastic integrity caused by heavy calcification burden. *Apoe^-/-^ Snrk^△SMC^* mice had a marked tendency for systemic calcification formation within lesion areas in the aortic root and brachiocephalic artery, compared with *Apoe^-/-^ Snrk^Flox^*mice (Figure 4C-4H, Figure S10). Thus, our findings suggest that SNRK deficiency aggravates vascular calcification formation *in vivo*.

**Figure 4.**
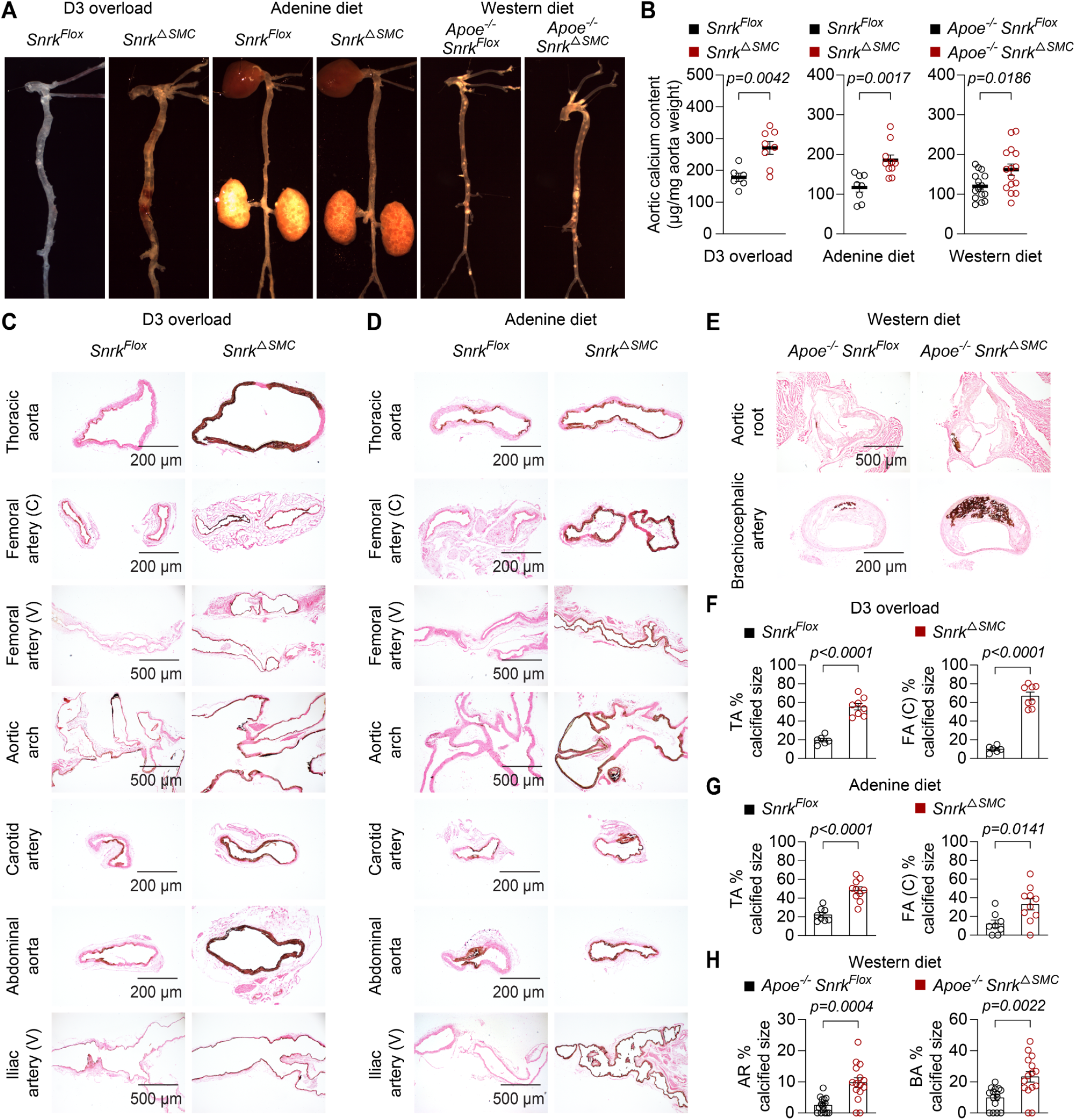
VSMC-specific SNRK deletion promotes vascular calcification in three mouse models of vascular calcification. (**A**) Representative en face view of aorta isolated from *Snrk^Flox^* or *Snrk^△SMC^* mice treated with either vitamin D3 or adenine diet, and *Apoe*^-/-^ *Snrk^Flox^* or *Apoe*^-/-^ *Snrk^△SMC^* mice fed with western diet (n = 6-15). (**B**) Aortic calcium content analysis of indicated mouse models of vascular calcification quantified by calcium assay and normalized to aortic dry weight, expressed as μg calcium per mg aorta (n = 6-15). (**C-H**) Representative von Kossa staining and calcification size analysis of thoracic aorta (TA), femoral artery (FA), aortic arch, carotid artery, abdominal aorta, iliac artery, aortic root (AR) and brachiocephalic artery (BA) of indicated mouse models of vascular calcification (n = 6-15). Cross section (C). Vertical section (V). Scale bars: 200 μm for thoracic aorta, femoral artery (C), carotid artery, and abdominal aorta images; 500 μm for femoral artery (V), aortic arch, and iliac artery (V) images for **C** and **D**. Scale bars: 500 μm for aortic root images; 200 μm for brachiocephalic artery images for **E**. Results are presented as mean ± SEM, and analyzed using two-tailed unpaired Student’s *t*-test for **D** and **F**, unpaired *t*-test with Welch’s correction for **G** and **H**.

### Calcification development in SNRK knockout mice is atherosclerosis-independent

To exclude the effect of atherogenesis on calcification development in *Apoe^-/-^ Snrk^△SMC^* mice, we next assessed absolute lesion areas and lesion area fractions in situ. We found no significant differences in both aortic root and brachiocephalic artery between the *Apoe^-/-^ Snrk^Flox^* and *Apoe^-/-^ Snrk^△SMC^* mice (Figure S11). *Apoe^-/-^ Snrk^Flox^* and *Apoe^-/-^ Snrk^△SMC^* mice had similar metabolic parameters, including serum triglyceride, cholesterol, blood glucose, and body weight (Figure S12). Thus, we conclude that the enhanced calcification formation in SNRK knockout mice is independent of atherosclerosis.

### SNRK depletion in VSMCs but not in endothelial cells and macrophages promotes vascular calcification *in vivo*

VSMCs, endothelial cells, and macrophages are central regulators of arterial calcification. We postulated that VSMCs, endothelial cells, and macrophages, which both express SNRK, participated in the SNRK-relevant modulation of calcification. Therefore, we also generated *Apoe^-/-^* background mice with a specific deletion of SNRK in endothelial cell (*Apoe^-/-^ Snrk^△EC^*, *Apoe^-/-^*-*Snrk^flox/flox^-Cdh5^Cre^*) or macrophage (*Apoe^-/-^ Snrk^△MΦ^*, *Apoe^-/-^*-*Snrk^flox/flox^-lysM^Cre^*), and fed them and littermates (*Apoe^-/-^ Snrk^Flox^*) with western diet for 24 weeks. Immunofluorescence staining revealed mild expression change of SNRK in endothelial cell and macrophage within brachiocephalic artery of *Apoe*^-/-^ mice fed with western diet for 24 weeks, compared with those fed with chow diet (Figure S13). Moreover, we found that SNRK deficiency in endothelial cell and macrophage had mild effect on calcification formation in situ in western diet-treated *Apoe^-/-^ Snrk^△EC^* mice and *Apoe^-/-^ Snrk^△MΦ^* mice (Figure S14). Taken together, these results indicate that VSMC-derived SNRK is crucial for regulating the progression of vascular calcification, but SNRK from endothelial cell and macrophage may not be involved.

### VSMC SNRK overexpression alleviates vascular calcification

To verify the beneficial effects of SNRK overexpression on calcification development, we isolated murine primary aortic VSMC and aortic rings from C57BL/6J mice. We incubated primary VSMC and aortic rings in osteogenic medium for 14 days, supplemented with adeno-associated virus (AAV) vector or AAV2-*Tagln*-*Snrk* (AAV-*Snrk*). Corresponding calcium nodules formation was detected by Alizarin red staining. Consistently, AAV-*Snrk*-mediated SNRK overexpression ameliorated calcium deposition in murine aortic VSMC (Figure 5A). It also resulted in reduced mineralization in aortic rings ex vivo (Figure 5B, Figure S15A). High phosphate exerted its deleterious effect on aggravation of osteogenic reprogramming by inducing RUNX2 and MSX2 expression in VSMC. The western blot results confirmed that SNRK overexpression suppressed the osteogenic factors induction, accompanied by restoring the loss of VSMC contractile marker α-SMA, comparing with the vehicle controls (Figure 5C). Moreover, we established an overexpression mouse model by injecting AAV-Vector or AAV-*Snrk* to *Snrk^Flox^* or *Snrk^△SMC^* mice 28 days prior to vitamin D3 supplement (Figure 5D). The efficiency of AAV-*Snrk* transfection was verified in VSMC and endothelium, as indicated in Figure S15B. As expected, vitamin D3 overload successfully triggered dilation and calcification of aortas, as evidenced by en face view of whole aorta morphology (Figure 5E), calcium content quantification (Figure 5F), and von Kossa staining in situ (Figure 5G). Less severe vascular calcification was confirmed by the alleviated aortic dilation, reduced calcium content, and decreased intensity of von Kossa staining both in the *Snrk^Flox^* and *Snrk^△SMC^* mice transfected with AAV-*Snrk* (Figure 5E-5G). Altogether, these results indicate that SNRK has a preventive effect on vascular calcification progression.

**Figure 5.**
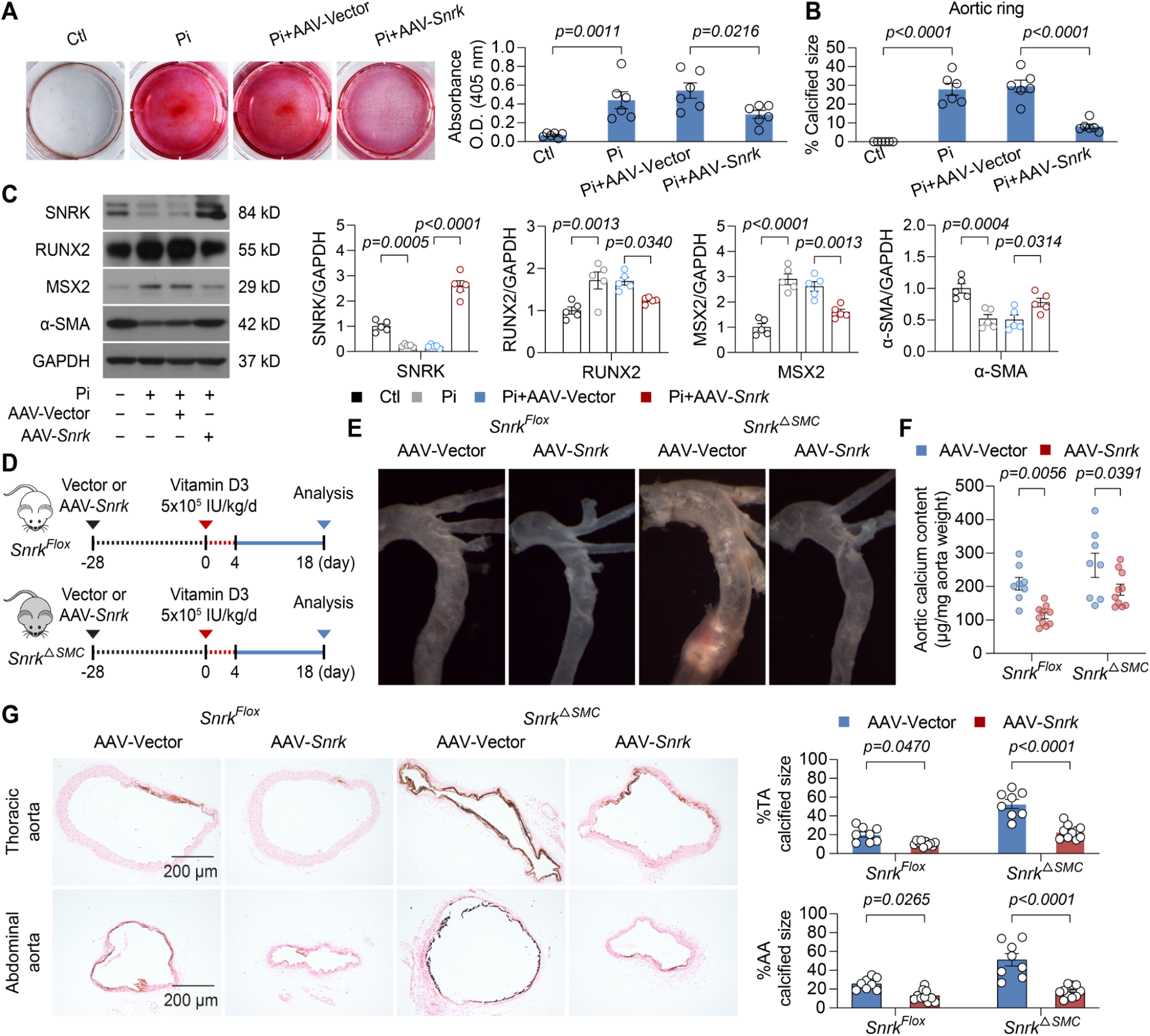
VSMC SNRK overexpression alleviates vascular calcification. (**A**) Representative whole-well alizarin red S staining of murine primary aortic VSMC isolated from C57BL/6J mice infected vector or AAV-*Snrk* treated with Pi for 14 days. Calcium deposition was quantified by measuring absorbance at 405 nm after dye extraction. Data are presented as relative calcium content (n = 6). (**B**) Calcification size analysis of aortic rings of C57BL/6J mice infected vector or AAV-*Snrk* treated with Pi for 14 days (n = 6). (**C**) Western blot analysis of SNRK and VSMC marker genes in Pi-treated murine primary aortic VSMC isolated from C57BL/6J mice infected vector or AAV-*Snrk* for 14 days (n = 5). (**D** and **E**) Experimental scheme and representative en face view of aorta isolated from *Snrk^Flox^* or *Snrk^△SMC^* mice transfected vector or AAV-*Snrk* 28-day prior to 4-day vitamin D3 overload. (**F**) Aortic calcium content analysis of indicated mice quantified by calcium assay and normalized to aortic dry weight, expressed as μg calcium per mg aorta (n = 8-10). (**G**) Representative von Kossa staining and calcification size analysis of thoracic aorta (TA) and abdominal aorta (AA) of indicated mice (n = 8-10). Scale bars: 200 μm. Results are presented as mean ± SEM, and analyzed using one-way ANOVA with Sidak’s post hoc test for **A-C**, two-way ANOVA with Sidak’s post hoc test for **F** and **G**.

### SNRK directly interacts with β-Catenin to modulate VSMC osteogenic reprogramming

To investigate the mechanism through which SNRK suppresses vascular calcification, we conducted an interactome analysis of SNRK in HASMCs transfected with Flag-SNRK-SBP. HASMCs transfected with the empty vector Flag-SBP were used as the negative control. Anti-SBP immunoprecipitates were analyzed by liquid chromatography-tandem mass spectrometry (LC-MS/MS). We then compared the 448 potential binding proteins identified by LC-MS/MS with the 257 vascular calcification-related proteins documented in DisGeNet, a database for human gene-disease associations (Figure 6A). The 11 potential binding proteins related to vascular calcification were β-Catenin (*CTNNB1*), glycogen synthase kinase 3β (*GSK3B*), glycogen synthase kinase 3α (*GSK3A*), cyclin-dependent kinase inhibitor 2A (*CDKN2A*), phospholipase D1 (*PLD1*), AMP-activated protein kinase subunit β1 (*PRKAB1*), calumenin (*CALU*), PDZ and LIM domain protein 7 (*PDLIM7*), reticulocalbin 2 (*RCN2*), cyclin T2 (*CCNT2*). Among them, β-Catenin, a central component of the Wnt signaling pathway, which governs osteoblast differentiation and vascular calcification, exhibited the highest association score (Figure 6A). Following coinfection with Flag-β-Catenin, GST, or GST-SNRK plasmids into HASMCs, β-Catenin was selectively immunoprecipitated by the anti-GST antibody in GST-SNRK group but not empty vector group (Figure 6B). We also demonstrated that endogenous SNRK associates with β-Catenin (Figure S16). To identify the specific binding motif of SNRK with β-Catenin, we subcloned 2 truncated SNRK N-terminal mutations with (aa 1-344) or without (aa 1-269) the ubiquitin-associated (UBA) domain (Figure 6C). HASMCs were transfected with GFP-β-Catenin and Flag-SBP, full-length Flag-SBP-SNRK, Flag-SBP-SNRK 1-269, or Flag-SBP-SNRK 1-350, followed by coimmunoprecipitation using the anti-SBP magnetic beads. As indicated in Figure 6D, full-length Flag-SBP-SNRK but not Flag-SBP-SNRK 1-269 or Flag-SBP-SNRK 1-350 efficiently immunoprecipitated with β-Catenin, suggesting that SNRK interacts with β-Catenin via its undefined C-terminal domain (aa 351-765). To elucidate more precisely how SNRK interacts with β-Catenin, we constructed the C-terminal domain aa 351-765 fragment (Flag-SBP-SNRK 351-765), a random aa 431-765 deletion mutant (Flag-SBP-SNRK 1-430), and the aa 270-765 fragment (Flag-SBP-SNRK 270-765) as shown in Figure 6C. The aa 270-765 fragment efficiently immunoprecipitated with β-Catenin, whereas the aa 351-765 fragment and aa 1-430 mutant construct did not (Figure 6D and 6E), suggesting the complete SNRK C-terminal structure together with UBA domain plays a crucial role in the overall binding with β-Catenin. Furthermore, confocal image of immunofluorescence staining showed that SNRK co-located with β-Catenin (Figure 6F). The active β-Catenin translocate to the nucleus to form a complex with T-cell factor/lymphoid enhancer factor (TCF/LEF), which drives the transcription of crucial osteogenic regulators such as RUNX2 and MSX2, contributing to vascular calcification. To investigate the role of SNRK on β-Catenin/TCF transcriptional activity, we transfected HASMCs with either TOPFlash reporter, responsive to TCF binding site, or the mutated FOPFlash reporter, serving as a control. Overexpression of SNRK, particularly the full-length form and the aa 270-765 fragment, resulted in decreased β-Catenin-stimulated TOPFlash/FOPFlash reporter activity. Conversely, neither the control Flag vector nor the aa 1-350 fragment affected the reporter activity (Figure 6G). These findings indicate that SNRK interacts with β-Catenin through aa 270-765 segment, which is the key fragment of the degradation domain.

**Figure 6.**
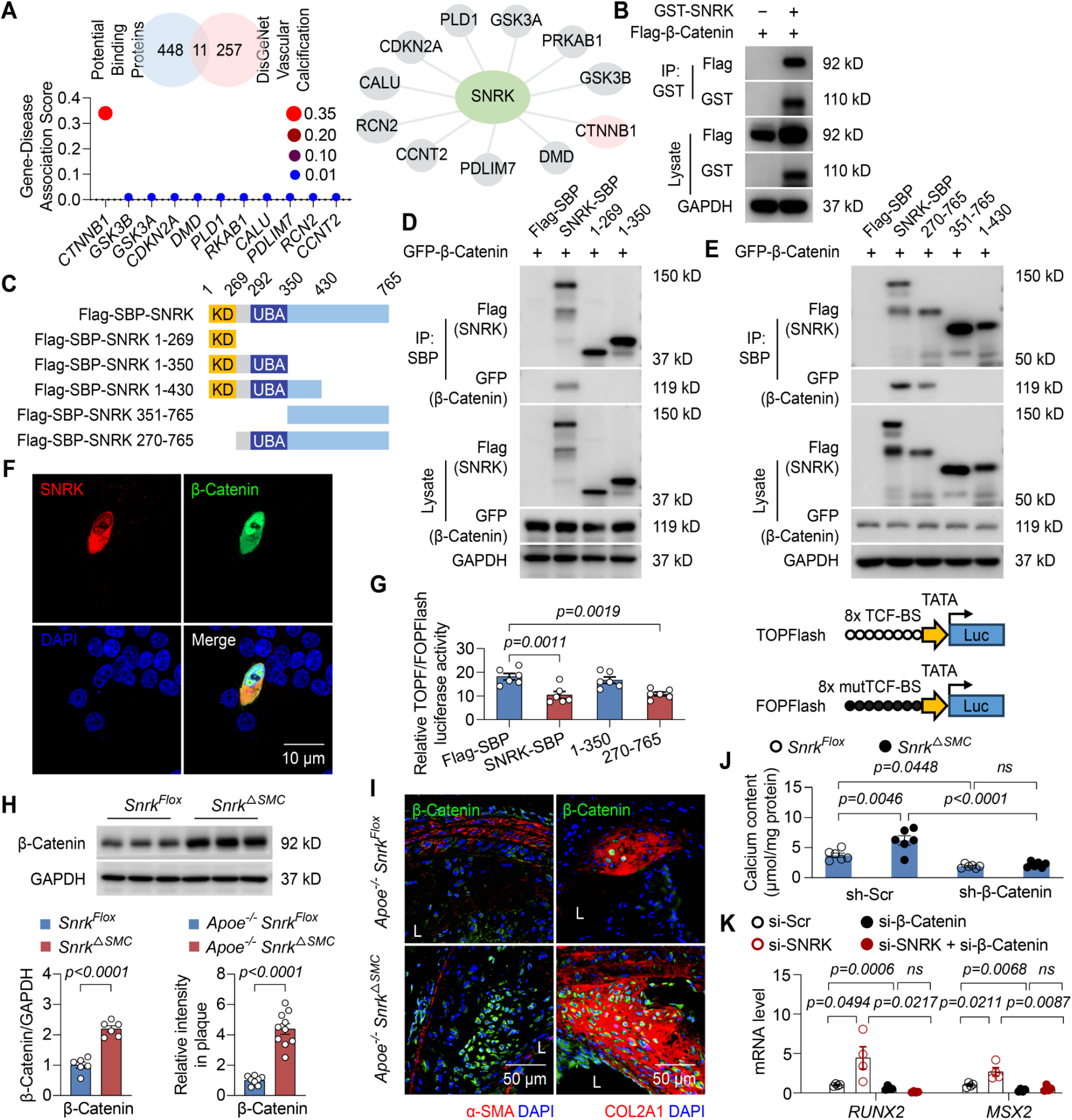
SNRK directly interacts with β-Catenin. (**A**) Comparison of potential binding proteins of SNRK derived from liquid chromatography-tandem mass spectrometry (LC-MS/MS) analysis with vascular calcification-related proteins in the database of human gene-disease associations (DisGeNet). Anti-Flag immunoprecipitates from HASMCs extracts transfected with Flag-SBP vector or Flag-SBP-SNRK for 48 hours cultured in osteogenic medium were analyzed by LC-MS/MS. (**B**) Co-IP assay of β-Catenin and SNRK in HASMCs transfected β-Catenin plasmid with or without SNRK plasmid for 48 hours (n = 4). (**C**) Schematic illustration of SNRK domains and SNRK serial deletion mutants. Kinase domain (KD), Ubiquitin-associated (UBA) domain. (**D** and **E**) Co-IP assay of SNRK serial deletion mutants in HASMCs co-transfected with β-Catenin (n = 4). (**F**) Representative confocal image of immunofluorescence staining of SNRK co-located with β-Catenin in HEK293T cells co-transfected with β-Catenin and SNRK (n = 5). Scale bar: 10 μm. (**G**) Schematic illustration and relative dual luciferase activity of HASMCs transfected with TOPFlash/FOPFlash, β-Catenin, Flag, full length SNRK plasmid or truncated SNRK plasmid (n = 6). (**H**) Western blot analysis of β-Catenin level in whole aorta of vitamin D3 treated *Snrk^Flox^* or *Snrk^△SMC^* mice (n = 6). (**I**) Quantification and immunofluorescence staining of β-Catenin co-located with α-SMA or COL2A1 in brachiocephalic artery of *Apoe*^-/-^ *Snrk^Flox^* or *Apoe*^-/-^ *Snrk^△SMC^* mice fed with western diet for 24 weeks (n = 8-10). Scale bar: 50 μm. (**J**) Calcium content quantification of aortic rings isolated from *Snrk^Flox^* or *Snrk ^△ SMC^* mice transfected with lentivirus of scramble (sh-Scr) or *Ctnnb1* shRNA cultured in osteogenic medium for 14 days (n = 6). (**K**) RT-PCR analysis of *RUNX2* and *MSX2* mRNA level in Pi-treated HASMCs transfected SNRK siRNA or/and β-Catenin siRNA for 24 hours (n = 4). Results are presented as mean ± SEM, and analyzed using two-tailed unpaired Student’s *t*-test for **H** and **I**, one-way ANOVA with Sidak’s post hoc test for **G**, and one-way ANOVA with Tukey’s post hoc test for **J** and **K**.

### SNRK modulates VSMC osteogenic reprogramming in a β-Catenin-dependent manner *ex vivo* and *in vitro*

Next, we detected β-Catenin level *in vivo*, result showed that the protein level of β-Catenin increased in whole aorta of vitamin D3 treated *Snrk^△SMC^* mice compared to the *Snrk^Flox^*mice (Figure 6H). Moreover, a significant increase in β-Catenin expression was observed by SNRK deficiency in the brachiocephalic artery of *Apoe*^-/-^ *Snrk^△SMC^* mice fed with western diet for 24 weeks, compared to the *Apoe*^-/-^ *Snrk^Flox^* mice (Figure 6I). To examine whether β-Catenin is involved in the vascular calcification promoted by SNRK deficiency, we transfected both scramble shRNA lentivirus particles or *Ctnnb1* shRNA in aortic rings isolated from *Snrk^Flox^* or *Snrk^△SMC^* mice cultured in osteogenic medium *ex vivo*. Indeed, aortic calcium deposition elevated in *Snrk^△ SMC^* mice, the additional β-Catenin knockdown effectively ameliorated this effect (Figure 6J). Similarly, to examine whether β-Catenin is involved in the VSMC osteogenic reprogramming promoted by SNRK deficiency, we transfected both SNRK and β-catenin siRNA in HASMCs. The silencing of β-Catenin eliminated the upregulation of RUNX2 and MSX2 mRNA and protein expression induced by SNRK deficiency (Figure 6K, Figure S17). Taken together, these results indicate that the inhibitory effect of SNRK on vascular calcification was mediated by β-Catenin.

### SNRK promotes β-Catenin ubiquitination and proteasomal degradation resulting in suppression of RUNX2 and MSX2 in VSMC

Next, we explored how the interaction between SNRK and β-Catenin inhibits β-Catenin signaling. Intriguingly, the mRNA levels of β-Catenin were not affected by silencing or overexpressing of SNRK in HASMCs (Figure 7A). This finding suggests that SNRK-repressed β-Catenin possesses a non-genomic role that extends beyond its function in transcriptional regulation. Reciprocally, the protein levels of β-Catenin were significantly induced in HASMCs by SNRK deficiency (Figure 7B). Since SNRK is a serine/threonine kinase, we investigated whether its kinase activity SNRK regulates β-Catenin, with SNRK activation via phosphorylation of the T-loop residue threonine-173. Our analysis involved the presence of either overexpression of wide-type SNRK or a kinase-dead SNRK mutant (SNRK-T173A), where threonine-173 in the catalytic domain was substituted with alanine. In both cases, the expression levels of β-Catenin, downstream RUNX2 and MSX2 were suppressed (Figure 7B, Figure S18). Taken together, these results suggest that the inhibition of β-Catenin by SNRK occurs independent of SNRK kinase activity. In addition, the overexpression of aa 1-350 fragment did not repress β-Catenin expression as the full-length SNRK and the aa 270-765 fragment (Figure 7C). To examine the mechanism by which SNRK repressed the protein level of β-Catenin, we assessed the stability of β-Catenin in the context of SNRK overexpression using protein synthesis inhibitor cycloheximide (CHX). The result showed that SNRK accelerated the instability and degradation of β-Catenin protein in the presence of CHX, compared with the vector control group (Figure 7D). Moreover, with proteasome inhibitor MG132 supplementation, the repression of β-Catenin was markedly prevented (Figure 7E). In accordance, full length SNRK and aa 270-765 SNRK fragment promoted the K48 ubiquitination of β-Catenin in VSMCs (Figure 7F and 7G), suggesting that SNRK enhanced β-Catenin degradation in a ubiquitin-proteasome-dependent pathway. The E3 ubiquitin ligases of the SCF complex (SKP1 [S-phase kinase associated protein 1]-Cullin1 (CUL1)-F-box protein) are responsible for the ubiquitination of β-Catenin. The F-box protein β-transducin repeat-containing protein (β-TrCP) acts as substrate of SCF complex to recognize β-Catenin for subsequent ubiquitination. Therefore, we verified the combination of SNRK, β-Catenin, and CUL1 complex. As depicted in Figure S19, endogenous binding of β-Catenin to SNRK, CUL1 but not CUL4A or CUL4B was evident, pulled down by β-Catenin immunoprecipitation. β-TrCP silencing alleviated the SNRK-induced β-Catenin reduction, and corresponding RUNX2 and MSX2 expression (Figure 7H, Figure S20A). In accordance, aa 270-765 SNRK domain were found to bridges endogenous β-TrCP and β-Catenin, coinciding with the formation of the β-Catenin–β-TrCP complex (Figure 7I, Figure S21). Next, we applied the SCF complex inhibitor MLN4924 in HASMCs together with SNRK overexpression, MLN4924 completely abolished SNRK-induced β-Catenin, RUNX2, and MSX2 repression (Figure 7J, Figure S20B). Moreover, SNRK overexpression markedly reduced β-Catenin protein levels and downstream RUNX2 and MSX2 expression in the presence of wild-type β-Catenin, whereas these effects were largely abolished when the β-TrCP degron-defective S33Y mutant was expressed (Figure S20C). In summary, SNRK promotes the ubiquitination and proteasomal degradation of β-Catenin in a β-TrCP degron-dependent manner.

**Figure 7.**
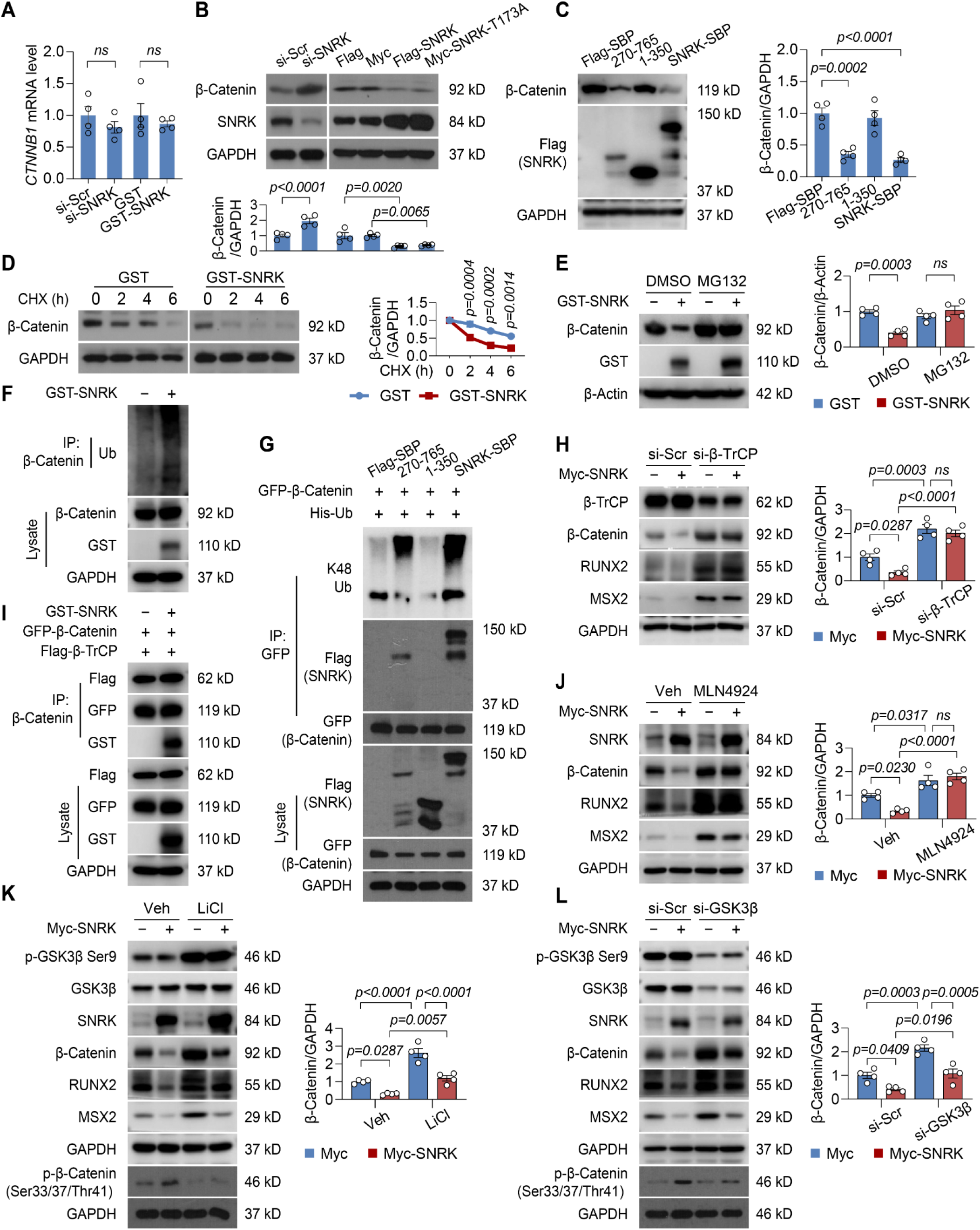
SNRK promotes β-Catenin ubiquitination and proteasomal degradation resulting in suppression of RUNX2/MSX2 in VSMC. (**A**) RT-PCR analysis of *CTNNB1* mRNA level in HASMCs transfected SNRK siRNA or SNRK plasmid for 24 hours (n = 4). (**B** and **C**) Western blot analysis of β-Catenin in HASMCs transfected SNRK siRNA, full length SNRK plasmid, SNRK kinase mutant (SNRK-T173A), or truncated SNRK plasmid as indicated for 48 hours (n = 4). (**D** and **E**) Western blot analysis of β-Catenin in HASMCs transfected SNRK plasmid with cycloheximide (CHX, 10 μg/mL) for different time, or MG132 (1 μmol/L) treatment for 8 hours (n = 4). (**F**) Co-IP assay of HASMCs transfected SNRK plasmid with 8-hour MG132 (1 μmol/L) pretreatment (n = 4). (**G**) Co-IP assay of HASMCs transfected β-Catenin, Ub, full length SNRK plasmid or truncated SNRK plasmid (n = 4). (**H**) Western blot analysis of β-Catenin in HASMCs transfected β-TrCP siRNA or SNRK plasmid as indicated (n = 4). (**I**) Co-IP assay of HASMCs transfected β-Catenin, β-TrCP, or SNRK plasmid (n = 4). (**J**) Western blot analysis of β-Catenin in HASMCs transfected SNRK and treated with vehicle or SCF complex inhibitor MLN4924 (1 μmol/L) for 6 hours (n = 4). (**K**) Western blot analysis of β-Catenin in HASMCs transfected SNRK and treated with vehicle or GSK3β inhibitor LiCl (20 mmol/L) for 24 hours (n = 4). (**L**) Western blot analysis of β-Catenin in HASMCs transfected GSK3β siRNA or SNRK plasmid as indicated (n = 4). Results are presented as mean ± SEM. *P* values are assessed using two-tailed unpaired Student’s *t*-test used for **A** and **B,** one-way ANOVA with Sidak’s post hoc test for **C** and **E-L**, Two-way ANOVA with Sidak’s post hoc test is used for **D**.

### SNRK promotes β-Catenin degradation through a mechanism independent of GSK3β destruction complex

In general, β-Catenin is inactivated by a destruction complex comprising GSK3β, adenomatous polyposis coli (APC), casein kinase 1 (CK1), and Axin. This protein complex mediates the phosphorylation of β-Catenin, which is then recognized, ubiquitinylated by SCF complex and subsequently degraded in proteasomes. Of note, GSK3β-Ser9 phosphorylation, which inactivates the kinase activity of GSK3β, was not affected by SNRK (Figure S22). LiCl, a direct inhibitor of GSK3β, did not affect SNRK repression of β-Catenin, RUNX2, and MSX2 (Figure 7K, Figure S20D). In accordance, GSK3β deficiency did not abolished the effect of SNRK on decreasing β-Catenin, RUNX2, and MSX2 expression in HASMCs. Moreover, pharmacological inhibition of GSK-3β by LiCl or genetic depletion of GSK-3β markedly lowers the basal phosphorylation of β-Catenin at Ser33/37/Thr41, but SNRK overexpression consistently modulated phosphorylation at these canonical degron residues and promoted β-Catenin degradation (Figure 7L, Figure S20E). Taken together, SNRK acts independently of GSK-3β kinase activity but converges on the degron-dependent degradation process.

## Discussion

In this study, we demonstrated that the deletion of SNRK in VSMCs, but not in endothelial cells or macrophages, significantly exacerbates osteogenic reprogramming of VSMCs and subsequent calcification. Importantly, overexpression of SNRK via AAV-*Snrk* infection effectively reversed this pathological phenotype in both *in vitro* and *in vivo* models. Our findings further revealed that the inhibitory effect of SNRK on vascular calcification is mediated through a β-Catenin-dependent mechanism (Figure 8). Specifically, SNRK binds directly to β-Catenin and facilitates the assembly of the β-TrCP and CUL1 E3 ubiquitin ligase complex. This interaction promotes the ubiquitin-mediated proteasomal degradation of β-Catenin, leading to its downregulation and suppression of osteogenic signaling pathways in VSMCs. Clinically, we observed that levels of PBMC *SNRK-AS1* were positively correlated with the progression of thoracic artery calcification in CKD patients. Taken together, our results suggest that by preventing osteogenic differentiation of VSMCs, SNRK acts as a safeguard against vascular calcification. These results not only advance our understanding of the molecular mechanisms underlying calcific vasculopathy but also suggest that SNRK may represent a promising therapeutic target for the prevention and treatment of vascular sclerosis in clinical contexts.

**Figure 8.**
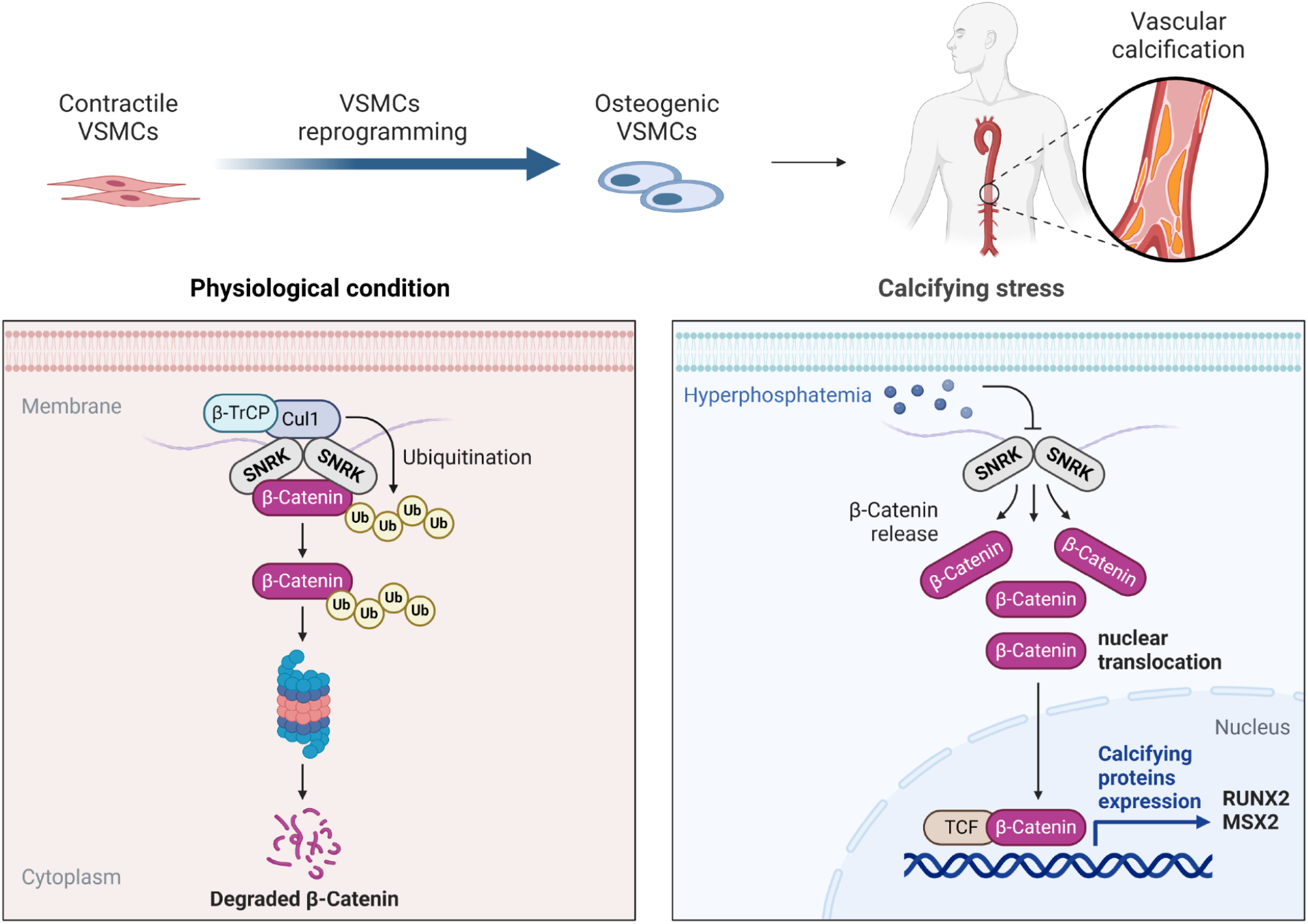
Graphical scheme of SNRK alleviating VSMCs osteogenic reprogramming and calcification by β-Catenin downregulation.

Emerging evidence suggests that circulating mononuclear cells participate in vascular calcification. In patients with subclinical atherosclerosis, PBMCs have been shown to express osteogenic markers osteocalcin, which correlate with arterial calcification burden.^36^ CD14+ monocytes interacted with the intermediate cells of the osteogenic SMC lineage by delivering TGFB1 and TNFSF10.^37^ These findings suggest that PBMCs reflect systemic pro-calcific vascular remodeling. Given the functional role of the *SNRK-AS1*/SNRK signaling axis in regulating VSMC calcification, our findings suggest that circulating *SNRK-AS1* levels may serve as a surrogate biomarker for vascular calcification severity.

Most importantly, current study identifies SNRK as a critical regulator of VSMC phenotype and vascular calcification by using three pathophysiological relevant mouse models. The vitamin D3 and adenine diet models mimic CKD-associated medial calcification driven by mineral imbalance, while the western diet model reflects chronic lipid- and inflammation-induced intimal calcification. Despite distinct initiating mechanisms, VSMCs are the predominant source of calcifying cells in both medial and intimal lesions, with lineage-tracing studies showing that 98% osteogenic-like cells in plaques originate from VSMCs.^38^ These three complementary models sharing common VSMCs osteogenic differentiation enabled us to assess whether SNRK uniformly regulates calcification across different pathological contexts. We consistently observed diminished expression of SNRK in the VSMC-rich medial layer of calcified vessels. This loss was accompanied by downregulation of VSMC contractile marker α-SMA and concomitant upregulation of osteogenic markers including RUNX2, MSX2, and OCN. These *in vivo* findings were corroborated by analysis of human femoral artery samples, in which SNRK expression was significantly lower in VSMCs within calcified regions compared to adjacent non-calcified areas. Together, our findings establish SNRK as a previously underappreciated but essential player in maintaining VSMCs contractile phenotype and suppressing their osteogenic differentiation, a central process in both medial and intimal vascular calcification.

Previous studies have implicated SNRK in vascular integrity and endothelial homeostasis. For instance, global heterozygous *Snrk* knockout mice exhibit impaired retinal angiogenesis and compromised neovascular responses following hindlimb ischemia.^34^ Similarly, endothelial-specific *Snrk* deletion replicates these vascular defects, and SNRK protects glomerular endothelial cells from angiotensin II-induced NF-κB/p65 pathway hyperactivation.^26^ These observations underscore the broad vascular relevance of SNRK. However, our study is the first to specifically dissect its functional role within VSMCs during vascular calcification.

To pinpoint the cellular contribution of SNRK, we employed a genetic loss-of-function strategy. Deletion of *Snrk* specifically in VSMCs resulted in exacerbated vascular calcification, whereas endothelial or macrophage-specific deletion produced only modest changes. This finding highlights a VSMC-autonomous role of SNRK in preventing calcific differentiation. Moreover, VSMC-specific *Snrk* deletion did not affect systemic metabolic parameters such as lipid profiles, glucose levels, or body weight, nor did it promote atherosclerotic plaque formation. This suggests that the pro-calcific phenotype arises directly from VSMC reprogramming, rather than from secondary metabolic alterations. Importantly, overexpression of SNRK in VSMCs effectively attenuated phosphate-induced calcific nodule formation in the aortic wall, providing direct functional evidence of its protective capacity. Therefore, these results strongly support the notion that SNRK acts as a molecular brake on VSMCs phenotypic switching and vascular mineralization.

Using LC-MS/MS-based proteomics with gene-disease association data analyses, we identified β-Catenin as a direct target of SNRK in regulating vascular calcification in current study. The pivotal role of the Wnt/β-Catenin signaling pathway in driving VSMC osteogenic differentiation is well-documented, as it directly targets key osteogenic factors such as RUNX2 and MSX2 under elevated phosphate stimuli in a bidirectional manner.^39-44^ Our findings revealed that SNRK deletion sustains elevated levels of β-Catenin in aortic calcified lesions, along with RUNX2 and MSX2. Clinical studies in dialysis patients indicate that circulating sclerostin levels, a Wnt inhibitor, are significantly correlated with the severity of vascular calcification.^45-47^ Although current therapies targeting the β-Catenin pathway mainly address cancer and osteoporosis, romosozumab remains the only approved agent that modulates Wnt/β-Catenin signaling to promote bone mineralization.^48,49^ Given the shared mechanisms between vascular calcification and bone formation, SNRK emerges as a potential therapeutic target by modulating β-Catenin pathways for mitigating vascular calcification.

Our study provides mechanistic insights into how SNRK acts as a negative regulator of β-Catenin. Post-translational modifications, particularly ubiquitination, tightly regulate β-Catenin stability, yet upstream modulators in vascular contexts are poorly defined.^50-53^ Under basal condition (Wnt-off), β-Catenin is phosphorylated by glycogen synthase kinase 3β (GSK3β) at Thr41, Ser37, and Ser33, promoting its recognition by β-TrCP for ubiquitination and proteasomal degradation. Upon Wnt activation (Wnt-on), β-Catenin dissociates this destruction complex, translocates into the nucleus, and activates gene transcription via TCF/LEF factors.^54^ SNRK has been implicated in β-Catenin regulation, with previous studies showing that SNRK promotes β-Catenin degradation in cardiomyocytes via GSK3β interaction,^55^ and reduces β-Catenin nuclear localization in colon cancer cells.^22^ In this study, we demonstrate that SNRK silencing increases β-Catenin protein levels without affecting its mRNA, indicating post-transcriptional regulation via the ubiquitin-proteasome pathway. β-TrCP, as part of the CUL1-containing ubiquitin ligase complex, targets β-Catenin and others for degradation,^56,57^ and our immunoprecipitation experiments confirmed SNRK’s interaction with this complex. Importantly, β-TrCP deficiency inhibited SNRK-mediated β-Catenin degradation and expression of downstream RUNX2 and MSX2, whereas GSK3β inactivation had no effect. Furthermore, we demonstrate that SNRK loss-induced calcification is β-Catenin-dependent. Collectively, these findings enhance understanding of noncanonical β-Catenin degradation in VSMCs and highlight SNRK activation as a potential strategy to counteract β-Catenin-driven vascular calcification.

Moreover, we found that SNRK facilitates β-Catenin degradation through its C-terminal region (aa 272-765), which contains the ubiquitin-associated (UBA) domain, consistent with previous reports of SNRK-mediated protein proteasomal degradation. However, the more specific domain underlying remains to be elucidated. Notably, SNRK overexpression reduced β-Catenin protein levels even in the absence of calcifying stimuli, suggesting that SNRK’s regulatory role on β-Catenin may extend beyond vascular calcification to broader intracellular processes governed by β-Catenin signaling, which merits further investigation.

Although the *Tagln* (SM22α) promoter is widely used to target VSMCs, SNRK expression is predominantly enriched in VSMC layer in our model, its activity can be induced in endothelial cells under angiogenic conditions as previously reported, indicating potential off-target activity in vascular models. Meanwhile, we did not use sh-β-Catenin–targeting virus as a direct *in vivo* rescue approach, which we acknowledge as a limitation for current study. We also acknowledge a sex limitation of our genetic model, as the *Myh11*-CreER^T2^ BAC transgene is inserted on the Y chromosome, only male offspring carrying Cre were included. Given sex differences in calcification pathophysiology, a well-powered assessment of specimens from both sexes is necessary for future study.

In summary, our findings establish SNRK as a critical suppressor of vascular calcification by inhibiting VSMC osteogenic differentiation. This study offers insights into future therapeutic strategies to prevent pathological cardiovascular calcification.

## Non-standard abbreviations and acronyms

ALP: Alkaline phosphatase
Apoe: Apolipoprotein E
COL1A1: Collagen I alpha1
COL2A1: Collagen II alpha1
CUL1: Cullin 1
ND: Normal diet
OCN: Osteocalcin
RUNX2: Runt-related transcription factor 2
α-SMA: Alpha smooth muscle actin
Ub: Ubiquitin
VSMC: Vascular smooth muscle cell
WD: Western diet

## Acknowledgments

None.

## Author contribution

M.H.Z. conceived and designed the research; L.O. and C.J.Y. performed the experiments and analyzed the data; H.H., X.Y.S., Z.Z.Z., C.J.K. and C.M.X. collected the human samples and clinical data; W.Y. and H.N. analyzed the transcriptomics data; Z.X.L. made critical revision of the manuscript; L.O. wrote the manuscript with comments from all authors; all authors approved the final version of the manuscript.

## Supplemental Materials

Expanded Methods

Figures S1 - S23

Tables S1 – S7

