## Supplemental data for "Sucrose Non-fermenting Related Kinase (SNRK) Deficiency-Mediated Osteogenic Differentiation in Vascular Smooth Muscle Cells Exacerbates Vascular Calcification"

**Running Title:** *SNRK in Vascular Calcification*

Liu Ouyang[1], MD, PhD, Chang-Jiang Yu[1,2], PhD, Wei-Dong Ye[3], PhD, Zhang Zheng-Zhipeng[4], MD, Huan Nie[5], PhD, Cheng Jin-Kun[4], MD, Xie Chen[4], PhD, Chang-Ming Xie[4], MD, Zhi-Xue Liu[1], PhD, Xiao-Yan Su[6], MD, Hui Huang[4], MD, PhD, Ming-Hui Zou[1,7], MD, PhD

1 Institute for Biomedical Sciences, Georgia State University, Atlanta, USA

2 Departments of Pharmacy and Cardiology, Harbin Medical University Cancer Hospital, Harbin, China

3 Department of Vascular Surgery, the Quzhou Affiliated Hospital of Wenzhou Medical University, Quzhou, China.

4 Department of Cardiology, the Eighth Affiliated Hospital of Sun Yat-sen University, Sun Yat-sen University, Shenzhen, China

5 Department of Social and Human Sciences, Institute of Science Tokyo, Tokyo, Japan

6 Department of Nephrology, Dongguan Tungwah Hospital, Dongguan, China

7 Department of Endocrinology and Metabolism, Tianjin Medical University General Hospital, Tianjin Medical University, Tianjin, China

**Authorship**: L.O., C.J.Y., and W.Y. contributed equally to this work.

**Address for Correspondence:**

Ming-Hui Zou, MD, PhD, 157 Decatur St SE, Atlanta, GA 30303, USA.

Hui Huang, MD, PhD, Shennan Middle Road, Shenzhen, 510275, China.

### Supplemental Methods

#### Study population

This case-sectional study was conducted at Tungwah Hospital, China, between March 2018 and January 2020. A total of 198 participants with chronic kidney disease (CKD) were initially enrolled as primary cohort. However, 10 participants were excluded from meeting the exclusion criteria, another 10 for missing peripheral blood collection, and 16 for lacking thoracic artery calcification data obtained through multidetector computed tomography (MDCT) scans. Written informed consent was obtained from all participants prior to their enrollment. For external validation, using the same inclusion and exclusion criteria, we collected an independent clinical cohort from another center between November 2023 and December 2025, the Eighth Affiliated Hospital of Sun Yat-sen University, consisting of a total of 110 patients with chronic kidney disease (CKD) from the Department of Nephrology. The study adhered to the ethical guidelines of the 1975 Declaration of Helsinki and received approval from the Ethics Committee of the Tungwah Hospital (2017DHLL017) and the Eighth Affiliated Hospital of Sun Yat-sen University (2023-100-01).

Chronic kidney disease (CKD) was diagnosed based on the 2012 Kidney Disease: Improving Global Outcomes (KDIGO) guidelines, which require evidence of a sustained decline in renal function lasting for more than three months.[1] Diagnostic criteria included an estimated glomerular filtration rate (eGFR) of 60 mL/min/1.73 $m^2$ or markers of kidney damage such as albuminuria (urine albumin ≥ 30 mg per 24 hours or urine albumin-to-creatinine ratio [ACR] ≥ 30 mg/g). The eGFR was calculated using the formula established in the Chinese version of Modification of Diet in Renal Disease (C-MDRD) study.[2] Participants were classified into CKD stages G1 - G2 (eGFR ≥60 mL/min/1.73 $m^2$ ) or G3 - G5 (eGFR <60 mL/min/1.73 $m^2$ ) based on kidney function prior to dialysis. Exclusion criteria for this study included the following: (1) age < 18 or ≥ 90 years; (2) pregnancy; (3) use of phosphate binders, vitamin D analogs, or corticosteroids within three months prior to recruitment; (4) acute renal failure; (5) uncontrolled hyperlipidemia; and (6) history of hepatic disease or neoplastic disease.

**Assessment of thoracic aorta calcification score**

Patients underwent a chest MDCT scan with standard electrocardiographically (ECG) gated protocol to evaluate thoracic aorta calcification. The thoracic aorta originates from the aortic root, consists of ascending aorta, aortic arch, and the descending aorta. The MDCT images were reconstructed with 1 mm slice thickness to assess vascular calcification. Agatston scores were calculated for the MDCT images, quantified by two independent technicians who were blinded to the clinical data, following standard protocols using Siemens Syngo CT Workplace software. In both asymptomatic and symptomatic patients, a thoracic artery calcification score of zero has been shown to independently predict a low risk of cardiovascular events or all-cause mortality over the medium and long term.[3,4] Therefore, the presence of calcification burden was defined as an Agatston score > 0 in the present study. All subjects are ethnical Chinese (Han), and the multiple linear regression analyses of clinical factors and thoracic artery calcification Agatston scores were adjusted for age and sex.

**Laboratory analyses**

Overnight fasting blood was collected from CKD patients, prior to dialysis, via median cubital vein for biochemical analysis. Peripheral blood mononuclear cells (PBMCs) were isolated using Histopaque-1077 (Sigma) density gradient. The sample was centrifuged at 400 × *g* for 20 minutes and the interface layer was collected as PBMCs. Genomic DNA was extracted from PBMCs using the DNeasy Blood & Tissue Kit from Qiagen. Agarose gel confirmed the integration of genomic DNA, and NanoDrop spectrophotometer measured the concentration. Plasma concentrations of potassium, sodium, chlorine, calcium, magnesium, phosphate, blood urea nitrogen (BUN), creatinine (CREA), uric acid (URCA), albumin (ALB), alanine transaminase (ALT), aspartate aminotransferase (AST), alkaline phosphatase (ALP), cholesterol (CHOL), triglyceride (TG), high-density lipoprotein cholesterol (HDL-C), and low-density lipoprotein cholesterol (LDL-C) were analyzed using the standard certified TBA-120 auto-analyzer (Toshiba Medical Systems) in the Central Clinical Laboratory at Donghua Hospital of Sun Yat-sen University. Clinical and biochemical parameters were collected from the patient electronic medical records in the hospital. The human autopsy femoral arteries samples (2101-00177) were purchased from National Disease Research Interchange (PA, USA).

**Animal experiments**

The animal experiments were conducted in accordance with protocols approved by the Georgia State University Institute Animal Care and Use Committee. Specific-pathogen-free mice were housed in the animal facility of Georgia State University under controlled conditions, including uniform temperature, humidity, and a 12-hour light-dark cycle. The mice had unrestricted access to sterilized water and the designated diet. *Snrk*$^{flox/flox}$ mice were kindly provided by Dr. Ramani Ramchandran at Medical College of Wisconsin. *Myh11-CreER*$^{T2+}$ (B6.FVB-Tg[Myh11-icre/ERT2]1Soff/J, Stock No. 019079), *Cdh5*$^{Cre}$ (B6.FVB-Tg[Cdh5-cre]7Mlia/J, Stock No. 006137), *LysM*$^{Cre}$ (B6.129P2-*Lyz2*$^{tm1(cre)Ifo}$/J, Stock No. 004781), and *Apoe*$^{-/-}$ (B6.129P2-*Apoe*$^{tm1Unc}$/J, Stock No. 002052) mice were purchased from the Jackson Laboratory (ME, USA). Of note, *Myh11-CreER*$^{T2+}$ has the *Myh11-CreER*$^{T2}$ BAC transgene inserted on the Y chromosome, thus only male offsprings will be used for experiments. *Snrk*$^{flox/flox}$ mice *(Snrk*$^{Flox}$) were crossed with *Myh11-CreER*$^{T2}$ mice to generate tamoxifen-inducible VSMC-specific SNRK knockout mice (*Snrk*$^{\triangle SMC}$) with C57BL/6J background. *Snrk*$^{flox/flox}$*-Myh11-CreER*$^{T2+}$ mice were further crossed with hyperlipidemic apolipoprotein E-deficient (*Apoe*$^{-/-}$) mice to generate *Apoe*$^{-/-}$ *Snrk*$^{\triangle SMC}$ (*Apoe*$^{-/-}$*-Snrk*$^{flox/flox}$*-Myh11-CreER*$^{T2+}$) mice and *Apoe*$^{-/-}$ *Snrk*$^{Flox}$ (*Apoe*$^{-/-}$*-Snrk*$^{flox/flox}$*-Myh11-CreER*$^{T2-}$) mice. Similarly, *Apoe*$^{-/-}$*-Snrk*$^{flox/flox}$ mice were crossbreed with *Cdh5*$^{Cre}$ or *LysM*$^{Cre}$ mice to generate mice targeted SNRK deletion in endothelial cell (*Apoe*$^{-/-}$ *Snrk*$^{\triangle EC}$, *Apoe*$^{-/-}$*-Snrk*$^{flox/flox}$*-Cdh5*$^{Cre}$) or macrophage (*Apoe*$^{-/-}$ *Snrk*$^{\Delta M\Phi}$, *Apoe*$^{-/-}$*-Snrk*$^{flox/flox}$*-LysM*$^{Cre}$).[5] *Snrk*$^{\triangle SMC}$ or *Apoe*$^{-/-}$ *Snrk*$^{\triangle SMC}$ mice aged 6-8 weeks were intraperitoneally injected with tamoxifen at 50 mg/kg body weight per mouse per day intraperitoneally for 5 consecutive days to specifically activate Cre recombinase in mature VSMC. Tamoxifen was dissolved in 20% ethanol/80% sunflower seed oil vehicle. Mice were monitored daily and used for further study 14 days after injection to ensure fully effective Cre recombinase.

For the vitamin D3 overload-induced mouse model of vascular calcification, mice were intraperitoneally injected with vitamin D3 at $5\times10^5$ IU/kg body weight per mouse per day for 4 consecutive days. Mice were monitored daily and terminated 14 days after injection. Vitamin D3 was dissolved in 4% ethanol/96% sunflower seed oil vehicle.[6]

For the adenine diet-induced CKD model, mice were randomly provided with regular chow diet for 8 weeks as the control group, or a special diet prepared by Research Diets Inc containing 0.2% adenine and 1.2% phosphorus as the CKD group.[7]

For the diet-induced vascular calcification model, mice were randomly provided with regular chow diet for 24 weeks as the control group, or diet that had a nutritional profile similar to western diet

(0.21% cholesterol, 40 kcal% fat, and 43 kcal% carbohydrate; D12079B, Research Diets Inc) for 24 weeks as the experimental group.[7]

Mice were fasted overnight and weighed before euthanasia, and blood was collected. The whole aorta from aortic root, aortic arch, brachiocephalic artery to femoral artery, was harvested and carefully dissected for further analyses. Fasting blood glucose levels were detected using an autoanalyzer. Serum levels of total cholesterol and triglyceride were measured using specific reagents (Infinity, Thermo Fisher Scientific).

**Chemicals and antibodies**

MG132 (M7449), cycloheximide (01810), DAPI (D9564), LiCl (L9650), and vitamin D3 (cholecalciferol, PHR1237) were obtained from Sigma-Aldrich (MO, USA). Penicillin-Streptomycin was from Thermo Fisher Scientific (MA, USA). Puromycin dihydrochloride (sc-108071) and polybrene (sc-134220) were purchased from Santa Cruz Biotechnology (TX, USA). MLN4924 (No. 15217) was from Cayman Chemical (MI, USA).

Antibodies against RUNX2 (12556S), ubiquitin (3936S), K48-linkage Specific Polyubiquitin (8081S), SRY-box transcription factor 9 (SOX9, 82630S), β-Catenin (8480S), Flag (14793S, 8146S), GST (2622S, 2624S), GFP (2956S, 2955S), Myc (2276S, 2278S), β-TrCP (4394S), p-GSK3β Ser9 (5558S), GSK3β (12456S), mouse IgG (5873S), and rabbit IgG (2729S) were purchased from Cell Signaling Technology (MA, USA). Antibodies against alpha smooth muscle actin (α-SMA, ab7817, ab5694) and vWF (ab11713) were purchased from Abcam (Cambridge, UK). Antibody against SNRK (NBP1-83668) was purchased from Novus Biologicals (CO, USA). The osteocalcin (OCN, sc-390877), msh homeobox 2 (MSX2, sc-365232), GAPDH (sc-32233), and β-Actin (sc-47778) antibody were purchased from Santa Cruz Biotechnology. The CD68

(MCA1957GA) was purchased from Bio-Rad (CO, USA). The collagen II alpha1 (COL2A1, MAB8887) was from Sigma-Aldrich (MO, USA).

**Luciferase reporter assay**

For the luciferase experiments, Human aortic VSMCs were transfected with β-Catenin, Flag or full length SNRK plasmid or truncated SNRK plasmid, as well as Super8×TOPFlash reporter (in which Firefly luciferase expression is driven through multiple TCF-binding sites upstream of a minimal TATA box; Addgene 12456) or Super8×FOPFlash reporter (in which the TCF-binding sites are mutated; Addgene 12457) plasmid, together with the pRL-TK vector expressing Renilla luciferase (1:10) to normalize for transfection efficiency. After 48 hours, the luciferase activity was measured using a Dual-Luciferase Assay Kit (E1910, Promega) according to the manufacturer's instructions. The value of the Firefly luciferase reporter gene activity was normalized to Renilla luciferase activity. The results were expressed as the ratio of TOPFlash over FOPFlash normalized luciferase activity.

**Immunofluorescence**

Cells grown on confocal culture plates or optimum cutting temperature (OCT) embedded cryosections (8 μm) were processed as previously reported.[7,8] Cells or slides were rinsed with deionized water. Slides were then fixed in 4°C acetone for 5 minutes, cells were fixed with 3.7% paraformaldehyde for 15 min at room temperature, then permeabilized with 0.2% Triton X-100 for 10 minutes, incubated with 5% goat serum (BioGenex, CA, USA) for 30 minutes to block non-specific binding. Slides were incubated with antibodies against SNRK, β-Catenin, RUNX2, vWF, CD68, COL2A1, or α-SMA at 4°C overnight, and then exposed to fluorescent label conjugated secondary antibodies (Alexa Fluor®, Invitrogen) for 1 hour at room temperature. Nuclei were counter-stained using DAPI for 10 minutes at room temperature. Images were captured using a microscope (IX73, OLYMPUS) or confocal microscopy (LSM 810, Carl Zeiss Microscopy Ltd, MA, USA), and were quantified using ImageJ (National Institutes of Health, USA). Results were expressed as the ratio of relative florescence intensity compared with the controls. All quantifications were performed by two observers without knowledge of the identities of samples. Human autopsy femoral arteries samples were embedded in paraffin. Sagittal sections were deparaffinized and rehydrated before staining with von Kossa solution for morphological analysis according to manufacturer's protocol.

**Calcification evaluation**

Calcified nodules were characterized using von Kossa stain (KTVKO, American MasterTech Scientific) and alizarin red stain (AA42746AP, Thermo Fisher Scientific). Briefly, aorta sections were immersed with 5% silver nitrate buffer and exposed to ultraviolet light for 30 minutes. After rinsing with deionized water, sections were incubated with 5% sodium thiosulfate, followed by staining with nuclear fast red stain. Calcium crystals appeared brown, cytoplasm was stained pink, nuclei were marked red. The percentage of calcification lesion size within the aortic medial layer were quantified using ImageJ.

VSMCs or aortic rings were cultured in osteogenic medium containing 2.6 mmol/L phosphate (Pi) to boost calcification formation. The aortic rings were dehydrated and embedded in paraffin, then sectioned into serial sections. The cells or aortic rings sagittal sections were incubated with 1% (w/v) Alizarin red S solution for 30 minutes following the manufacturer's protocol. The red color indicated a positive stain result. Then the alizarin red S staining solution was removed, 10% acetic acid was added to stained cells to incubate for 30 minutes at room temperature. Ammonium hydroxide (10%; Sigma-Aldrich) was added to before measuring the absorbance of solubilized calcium-bound alizarin red S at 405 nm using TECAN Spark microplate reader. The percentage of calcification lesion size within the aortic ring were quantified using ImageJ.

Cell lysates or homogenized aortic tissues were treated with 0.6 mol/L HCl overnight at 4°C. Released calcium in the content in the supernatant was quantified using the Calcium Assay Kit (ab102505, Abcam) in compliance with the manufacturer’s protocol. Calcium content was calibrated by the weight of aorta or protein concentration.

**Cell and aortic ring culture**

Primary VSMCs were isolated from aortas of mice aged 6-8 weeks and maintained in 1640 medium (Lonza, USA) supplemented with 15% (v/v) fetal bovine serum (FBS, GIBCO, USA) at 37 °C, under a humidified environment with 5% CO2 and 95% air as previously described.[9] Human aortic VSMCs were obtained from the American Type Culture Collection (ATCC, MD, USA), and cultured in M231 medium (GIBCO, USA) containing 5% FBS, with smooth muscle growth supplement, 100 U/mL penicillin, and 100 μg/mL streptomycin. HEK293T cells were purchased from ATCC and maintained in Dulbecco's modified eagle's medium (DMEM, GIBCO, USA) containing 5% FBS. HEK293T cells or VSMCs (passage numbers 3 to 6) reaching 70-80% confluence were used for the experiments. Thoracic aortas were isolated from mice, dissected to remove adventitia under sterile conditions. The aortas were then sectioned into 5-mm rings and cultured in DMEM containing 10% FBS.

**siRNA transfection**

Small interfering RNAs (siRNAs) targeting β-Catenin (sc-29209), β-TrCP (sc-37178), or GSK3β (sc-35527) were purchased from Santa Cruz Biotechnology. The siRNA mixture targeting SNRK (AM51331) and SNRK antisense RNA 1 (*SNRK-AS1*) was obtained from Thermo Fisher Scientific

(MA, USA). Cells were transfected siRNA (20 nmol/L) together with Lipofectamine RNAiMAX Reagent (13778150, Invitrogen), in accordance with the manufacturer's protocol.

**Lentivirus construction and transfection**

Lentivirus expressing *SNRK* shRNA was packaged in HEK293T cells using SNRK shRNA (sc-78359-SH) plasmid, pMD2.G (12259, Addgene) plasmid and psPAX2 (12260, Addgene) using polyethylenimine. Supernatant was harvested 48 hours post-transfection and filtered with a 0.45 μm PVDF filter, and then virus was concentrated using Lentivirus purification kit (G171, ABM), and lentiviral titers were measured by qPCR Lentivirus Titration kit (LV900, ABM). Cells were infected with recombinant lentiviral particles carrying control short hairpin RNA (shRNA) (sc-108080, Santa Cruz Biotechnology), β-Catenin shRNA (sc-29210-V, Santa Cruz Biotechnology) or SNRK shRNA at a multiplicity of infection (MOI) of 10, together with polybrene (8 μg/mL), according to the manufacturer's instructions. Stable colonies were selected by 5 μg/mL puromycin dihydrochloride for 7 days.

**Plasmid construction and transfection**

The human *SNRK* (NM_017719) gene was amplified from pLenti-C-mGFP-P2A-Puro-SNRK (sc114000, Origene) using RT-PCR, the flanking endonuclease SalI/NotI restriction sites were designed into the PCR primers. Then PCR product and pHAGE-Flag-SBP vector were digested with SalI (R3138L, NEB) and NotI (R3189L, NEB) prior to ligation with T4 DNA ligase (M0202L, NEB), resulting in a mammalian expression construct for SNRK (Flag-SBP-SNRK). The entire insert in the SNRK expression construct was sequenced and verified. In accordance, the cDNA fragments that encoded SNRK aa 1-269, SNRK aa 1-350, SNRK aa 1-430, SNRK aa 351-765, and SNRK aa 270-765 were amplified using specific designed primers. GST-SNRK, Myc-SNRK, Flag-SNRK were generated using empty vectors with indicated tag. The mutant for the kinase inactive SNRK-T173A (Myc-SNRK-T173A) was generated using previously established pCMV-SNRK-T173A-HA plasmid (31086184). The primers were listed in Supplementary Table 7. pEGFP-N1-β-Catenin (71367), pcDNA3-Flag-β-Catenin (16828), pcDNA3-S33Y β-Catenin (19286), pcDNA3-myc3-CUL1 (19896), pcDNA3-myc3-CUL4B (19922), and pcDNA3-myc3-CUL4A (19951) were purchased from Addgene (MA, USA). Plasmids were transfected into VSMCs by electroporation using the P1 Primary Cell 4D-Nucleofector X Kit L from Lonza (MD, USA) following the manufacturer's instructions. Plasmids were transfected into HEK293T cells using lipofectamine transfection reagent 2000 (11668-019, Invitrogen) following the manufacturer's instructions.

**Recombinant adeno-associated virus construction and infection**

To generate adeno-associated virus (AAV) encoding mouse *Snrk* (NM_133741), we transfected the genomic AAV vector ligated with *Tagln* promoter (Figure S23), AAV helper plasmid, and AAV-2 plasmid into HEK293 cells, then harvested and purified viral particles as previously described.[10] RT-PCT was performed to amplify the *Snrk* open reading frame clone from pLenti-C-Myc-DDK-P2A-Puro-*Snrk* plasmid (Origene, MR217507). The primers for RT-PCR are listed in Supplemental Table 7. Mice (~20 g) received 100 μL AAV2-*Tagln*-*Snrk* via tail vein, corresponding to $2 \times 10^{11}$ genome copies (GC) per mouse (~$1 \times 10^{13}$ GC/kg; viral titer $2 \times 10^{12}$ GC/mL), as previously described.[11]

**Immunoprecipitation and western blot analysis**

For immunoprecipitation, cells were treated with MG132 (1 μmol/L) for 8 hours to inhibit proteasomal degradation prior to being lysed in non-denaturing buffer (20 mmol/L Tris-HCl pH 7.5, 150 mmol/L NaCl, 10% glycerol, 1% Triton X-100) containing protease inhibitor cocktail (5871S, Cell Signaling Technology). The lysates were incubated with targeted primary antibody overnight, followed by incubation with IgG magnetic beads (S1431S, NEB) for 2 hours at 4°C with gentle rotation. The beads were resuspended in 2× Laemmli buffer (BP-111NR, Boston Bioproducts) and briefly boiled at 99°C for complete denaturation before immunoblotting. For streptavidin binding peptide (SBP)-tagged proteins pull-down, the supernatant was directly incubated with streptavidin magnetic beads (NEB, S1420S) overnight at 4°C with gentle rotation before denaturation.

The endothelium of aorta sample was mechanically removed using a cotton swab, and the adventitia was carefully dissected. Approximately 200 μg of total protein was obtained from a single aortic segment (aortic arch to femoral artery) per mouse. Cellular lysates or aortic tissue homogenates were prepared on ice using RIPA lysis buffer (sc-24948A, Santa Cruz Biotechnology) for subsequent western blot analysis. The protein concentrations were assessed by Pierce BCA Protein Assay Kit (23225, Thermo Fisher Scientific). Equivalent amounts of protein (20 - 40 μg) were separated by SDS-PAGE and then electrophoretically transferred onto nitrocellulose membranes. The membranes were incubated with primary antibody overnight at 4°C, followed by incubation with horseradish peroxidase-conjugated secondary antibody at room temperature for 1 hour. Western blots and dot plots represent independent biological replicates. The integrated intensity of individual protein band was quantified based on density using ImageJ software.

**SNRK interactome**

SNRK-interacted proteins were identified by liquid chromatography tandem mass spectrometry (LC-MS/MS) as previously described.[12] In summary, HASMC were transfected with either Flag-SBP vector (negative control) or Flag-SBP-SNRK for 48 hours. The harvested cell lysates were immunoprecipitated with streptavidin magnetic beads overnight at 4°C. Ten percent of the beads were taken out to boil with SDS loading/sample buffer to elute proteins for analysis. Western blot and Coomassie-stained gel were used to confirm the presence of the target protein and evaluate the protein amount yield from the pull-down. The remaining proteins on beads were eluted, alkylated, digested with trypsin, and subjected to LC-MS/MS. Specific SNRK-associated proteins were compared with vascular calcification-related proteins from the DisGeNet human gene-disease association database.[13] Among the identified potential binding proteins, the protein with the highest association score in the database, related to vascular calcification, was selected for further analysis.

**Quantitative real time polymerase chain reaction (RT-PCR)**

Total RNA was extracted from VSMCs using TRIzol reagent (15596-026, Invitrogen), and reverse transcription was carried out using the iScript cDNA Synthesis Kit (1708890, Bio-Rad). PCR amplification was performed in triplicate with iQ SYBR Green supermix, utilizing the CFX96 Touch real-time PCR system (Bio-Rad, USA). The primers for RT-PCR are listed in Supplemental Table 7. The relative cycle threshold values of genes were calculated using the $2^{-\Delta\Delta CT}$ method, with normalization to the expression levels of housekeeping gene.

**RNA-seq processing and differential expression analysis**

Bulk RNA-seq datasets of C57BL/6 mouse aortic tissues, including healthy and calcified samples, were retrieved from the Gene Expression Omnibus (GEO). To harmonize data processing across studies, raw FASTQ files were downloaded and reanalyzed using a unified pipeline. Reference indices for STAR (v2.7.11b) and Salmon (v1.10.1) were prepared based on the mouse reference genome GRCm39 with GENCODE vM38 annotation. Adapter trimming and read-level quality control were performed using fastp (v0.23.4). Clean reads were aligned to GRCm39 with STAR, and sequencing quality was assessed with FastQC (v0.12.1). Gene-level quantification was conducted using featureCounts (Subread v2.0.6) to generate a raw count matrix. Given that most datasets were derived from poly(A)-enriched mRNA libraries, only protein-coding and lncRNA genes annotated in GENCODE were retained for downstream analysis. Low-expression genes were filtered using filterByExpr in edgeR. To correct batch effects while preserving group-associated biological differences, ComBat_seq was applied to the filtered raw counts. Differential expression analysis was then performed using DESeq2 with the design formula - group, with healthy aorta samples set as the reference level.

**Bioinformatics Analysis of Single-Cell Transcriptome Sequencing Data**

We used "calcified atherosclerotic plaque" and "single-cell RNA sequencing" as keywords to search and screen suitable datasets from the Gene Expression Omnibus (GEO) database, and finally identified the GSE159677 dataset for subsequent analysis. The GSE159677 dataset contains the single-cell transcriptome of entire calcified atherosclerotic core (AC) plaques and patient-matched proximal adjacent (PA) portions of carotid artery tissue from three patients undergoing carotid endarterectomy. The GSE159677 dataset was imported into Seurat (version 5.2.1), and quality control and downstream analyses were performed following standard protocols. For quality control, cells were retained if they met the following criteria: expressing 200–5000 genes per cell, having fewer than 20,000 Unique Molecular Identifiers (UMIs) per cell, and with mitochondrial gene UMI proportion less than 10%. To correct batch effects among different samples, the Canonical Correlation Analysis (CCA) model was used for data integration. Specifically, the "FindIntegrationAnchors" and "IntegrateData" functions in the Seurat package were employed to identify "anchors" between datasets, thereby achieving integration of data from different samples. Subsequently, data dimensionality was reduced to the top 30 Principal Component Analysis (PCA) components via principal component analysis. Cell clusters were identified using the "FindClusters" function with a resolution of 0.5. Uniform Manifold

Approximation and Projection (UMAP) dimensionality reduction technology was adopted for data visualization. Differentially expressed genes (DEGs) were identified using the "FindMarkers" function. Cell type annotation was performed based on previously reported marker genes in the literature.[14,15] GO and KEGG enrichment analyses were conducted using the R package clusterProfiler (version 4.14.0), with a corrected $p < 0.05$ considered significantly enriched.

**Statistical analyses**

All experimental results were expressed as mean ± standard error of the mean (SEM), and clinical results as mean ± standard deviation (SD), median (25th-75th quartiles) for continuous variables, or n (%) for categorical variables, respectively. The Shapiro-Wilk test or D'Agostino-Pearson test was performed to assess data normality. The equality of group variances was assessed using *F* test or Brown-Forsythe test. For comparisons between two groups, significance was assessed using unpaired two-tailed Student's *t*-tests, unpaired *t*-test with Welch's correction, or nonparametric Mann-Whitney *U* test. For comparisons among multiple groups, analysis of variance (ANOVA) or Welch's ANOVA test accompanied by Tukey's post hoc test (equal variances assumed), Dunnett's post hoc test, or Sidak post hoc test (equal variances not assumed) was performed. Correlation was assessed using Spearman's rank correlation coefficient. Model discrimination was assessed using the c-Statistic (AUC), with differences between models evaluated by the DeLong test. Reclassification was assessed using integrated discrimination improvement (IDI) and continuous net reclassification improvement (NRI). Missing values were imputed using multiple imputations. Analyses were subsequently conducted on the imputed datasets. All statistical analyses were performed by two independent researchers in a blinded manner. $P < 0.05$ was considered statistically significant. Statistical analyses were performed using SPSS version 29.0 (IL, USA) or GraphPad Prism 9.5 (CA, USA).

**Supplemental Tables**

**Supplemental Table 1.** Basic characteristics in CKD patients from primary cohort (Tungwah Hospital) with or without thoracic artery calcification

| Characteristics | Non-VC (n = 77) | VC (n = 85) | *p* value |
|---|---|---|---|
| **Demographic characteristics** | | | |
| Male, n (%) | 50 (64.9%) | 52 (61.2%) | 0.622 |
| Age, years | 57 (48 - 69) | 69 (60 - 76) | 0.000 |
| BMI, $kg/m^2$ | 23.44 (20.83 - 25.95) | 23.11 (20.53 - 25.90) | 0.441 |
| SBP, mmHg | 138.6 ± 23.33 | 146 ± 20.8 | 0.034 |
| DBP, mmHg | 84 (76 - 93) | 79 (69 - 86) | 0.001 |
| **Plasma biochemical characteristics** | | | |
| K, mmol/L | 4.18 ± 0.66 | 4.39 ± 0.64 | 0.050 |
| Na, mmol/L | 139 (137 - 141) | 138 (135.55 - 141) | 0.049 |
| Cl, mmol/L | 101 (97.05 - 105.3) | 98.7 (93.55 - 102.3) | 0.004 |
| Ca, mmol/L | 2.15 (2.03 - 2.25) | 2.18 (2.075 - 2.29) | 0.371 |
| Mg, mmol/L | 0.84 ± 0.15 | 0.85 ± 0.16 | 0.604 |
| Pi, mmol/L | 1.25 (0.955 - 1.62) | 1.31 (1.065 - 1.76) | 0.291 |
| GLU, mmol/L | 5.12 (4.34 - 6.01) | 5.51 (4.62 - 7.35) | 0.072 |
| BUN, mmol/L | 12.34 (5.59 - 21.21) | 16.68 (9.8 - 22.61) | 0.043 |
| CREA, μmol/L | 458 (116.45 - 1043.1) | 552.7 (247.9 - 926.5) | 0.255 |
| CysC, mg/L | 12.34 (4.56 - 56.75) | 8.24 (4.54 - 18.65) | 0.133 |
| eGFR, mL/min·per $1.73m^2$ | 4.46 (2.38 - 6.20) | 4.31 (1.35 - 6.56) | 0.368 |
| URCA, μmol/L | 411.48 ± 131.25 | 379.74 ± 128.64 | 0.122 |
| PALB, mg/dL | 26.78 ± 7.76 | 27.83 ± 8.96 | 0.429 |
| TP, g/L | 67.03 ± 7.3 | 65.74 ± 10.3 | 0.354 |
| ALB, g/L | 36.67 ± 6.52 | 34.62 ± 4.69 | 0.025 |
| GLOB, g/L | 30.6 (27.25 - 34.2) | 29.4 (25.25 - 33.85) | 0.288 |
| A/G | 1.16 (1.02 - 1.33) | 1.2 (0.99 - 1.4) | 0.742 |
| TBIL, μmol/L | 6.3 (4.65 - 7.55) | 6 (4.4 - 9.15) | 0.829 |
| DBIL, μmol/L | 2.4 (1.75 - 3.35) | 2.4 (1.7 - 3.15) | 0.757 |
| ALT, U/L | 13 (8 - 22.5) | 10 (7 - 16) | 0.021 |
| AST, U/L | 16 (13 - 22) | 15 (12 - 20) | 0.214 |
| ALP, U/L | 70 (51.5 - 83.5) | 79 (60.5 - 106.5) | 0.006 |
| GGT, U/L | 25 (15 - 40.5) | 24 (15.5 - 39.5) | 0.754 |
| LDH, U/L | 202 (169.5 - 237.5) | 221 (177.5 - 263) | 0.100 |
| TBA, μmol/L | 4 (2 - 7) | 3 (2 - 6) | 0.227 |
| CHE, U/L | 5555 (4353.5 - 6759.5) | 6086 (4835.5 - 7320) | 0.065 |
| CK, U/L | 68 (43.5 - 138.5) | 79 (48.5 - 137) | 0.526 |
| CKMB, U/L | 16 (12 - 20) | 14 (10 - 18.5) | 0.179 |
| HBDH, U/L | 166 (135.5 - 194.5) | 181 (145.5 - 228.5) | 0.065 |
| ApoA, g/L | 1.2 ± 0.33 | 1.15 ± 0.32 | 0.309 |

| | | | |
|---|---|---|---|
| ApoB, g/L | 0.85 ± 0.29 | 0.87 ± 0.32 | 0.661 |
| CHOL, mmol/L | 4.25 ± 1.29 | 4.02 ± 1.1 | 0.216 |
| TG, mmol/L | 1.5 (0.9 - 2.3) | 1.28 (0.93 - 1.92) | 0.163 |
| HDLC, mmol/L | 1.05 (0.79 - 1.38) | 0.99 (0.74 - 1.36) | 0.586 |
| LDLC, mmol/L | 2.6 ± 1.2 | 2.46 ± 0.94 | 0.408 |
| *SNRK-AS1* | 0.95 (0.75 - 1.26) | 1.52 (1.21 - 1.90) | 0.000 |

Values are expressed as mean ± SD or median (25th to 75th quartiles) for continuous variables and n (%) for categorical variables, respectively. Statistical significance was assessed using Student's t test or nonparametric Mann Whitney U test. BMI, body mass index; SBP, systolic blood pressure; DBP, diastolic blood pressure; K, potassium; Na, sodium; Cl, chlorine; Ca, calcium; Mg, magnesium; Pi, phosphate; GLU, glucose; BUN, blood urea nitrogen; CREA, creatinine; CysC, Cystatin C; eGFR, estimated glomerular filtration rate; URCA, uric acid; PALB, prealbumin; TP, total protein; ALB, albumin; GLOB, globulin; A/G, albumin/globulin ratio; TBIL, total bilirubin; DBIL, direct bilirubin; ALT, alanine transaminase; AST, aspartate aminotransferase; ALP, alkaline phosphatase; GGT, gamma glutamyl transferase; LDH, lactate dehydrogenase; TBA, total bile acids; CHE, cholinesterase; CK, creatine kinase; CKMB, Creatine Kinase MB; HBDH, hydroxybutyrate dehydrogenase; ApoA, apolipoprotein A; ApoB, apolipoprotein B; CHOL, total cholesterol; TG, triglyceride; HDLC, high-density lipoprotein cholesterol; LDLC, low-density lipoprotein cholesterol.

**Supplemental Table 2.** Multiple linear regression analysis of association of *SNRK-AS1* and clinical factors and thoracic artery calcification Agatston Score adjusted for age and sex in primary cohort

| Variable | Coefficient (B) | 95% CI | | | *p* value |
|---|---|---|---|---|---|
| Gender | -1.086 | -7.948 | to | 5.777 | 0.755 |
| Age, years | 0.489 | 0.254 | to | 0.724 | **<0.001** |
| BMI, kg/m$^2$ | 0.819 | -0.081 | to | 1.719 | 0.074 |
| SBP, mmHg | 0.121 | -0.029 | to | 0.272 | 0.113 |
| Pi, mmol/L | 9.617 | 3.108 | to | 16.126 | **0.004** |
| GLU, mmol/L | 1.121 | -0.201 | to | 2.444 | 0.096 |
| eGFR, mL/min·per 1.73m$^2$ | -0.656 | -1.647 | to | 0.336 | 0.193 |
| URCA, μmol/L | 0.010 | -0.017 | to | 0.037 | 0.459 |
| ALB, g/L | -0.532 | -1.144 | to | 0.079 | 0.087 |
| ALT, U/L | -0.011 | -0.067 | to | 0.046 | 0.704 |
| ALP, U/L | 0.076 | 0.038 | to | 0.115 | **<0.001** |
| TG, mmol/L | 0.895 | -0.747 | to | 2.538 | 0.283 |
| HDLC, mmol/L | 1.865 | -6.372 | to | 10.102 | 0.655 |
| LDLC, mmol/L | -0.653 | -3.748 | to | 2.442 | 0.677 |
| *SNRK-AS1* | 10.061 | 6.950 | to | 13.173 | **<0.001** |

Abbreviations are defined as in Supplemental Table 1. In consideration of the variation among clinical variables, the value of thoracic artery calcification scores were taken a square root transformation for analysis, and the regression residuals are normally distributed.

**Supplemental Table 3.** Incremental predictive performance after adding *SNRK-AS1* to the model containing conventional risk factors in primary cohort

| Variable | c-Statistics AUC (95% CI) | ΔAUC (95% CI) | *p* value | NRI | IDI (95% CI) | *p* value |
|---|---|---|---|---|---|---|
| Conventional risk factors | 0.812 (0.748-0.877) | | | | | |
| Conventional risk factors +*SNRK-AS1* | 0.927 (0.891-0.964) | 0.115 (0.061-0.169) | <0.001 | 1.21 | 0.258 (0.190 - 0.327) | <0.001 |

NRI, Net Reclassification Improvement; IDI, integrated discrimination improvement; The baseline model included vascular calcification conventional risk factors including sex, age, BMI, SBP, Pi, GLU, eGFR, URCA, ALB, ALT, ALP, TG, HDL-C, LDL-C. Abbreviations are defined as in Supplemental Table 1.

**Supplemental Table 4.** Basic characteristics of CKD patients from the external validation cohort (Eighth Affiliated Hospital of Sun Yat-sen University) stratified by median thoracic artery calcification (TAC) score

| Characteristics | Lower TAC (n = 55) | Higher TAC (n = 55) | *p* value |
|---|---|---|---|
| **Demographic characteristics** | | | |
| Male, n (%) | 32 (58.2%) | 33 (60.0%) | 0.459 |
| Age, years | 50 ± 13 | 56 ± 14 | 0.015 |
| SBP, mmHg | 142 ± 27 | 146 ± 23 | 0.519 |
| DBP, mmHg | 92 ± 16 | 88 ± 17 | 0.157 |
| **Plasma biochemical characteristics** | | | |
| K, mmol/L | 4.51 (4.16 - 5.15) | 4.90 (4.51 - 5.35) | 0.144 |
| Na, mmol/L | 138.21 (135.23 - 140.24) | 137.47 (135.64 - 139.30) | 0.095 |
| Cl, mmol/L | 101.15 ± 4.69 | 100.60 ± 4.21 | 0.105 |
| Ca, mmol/L | 2.24 ± 0.16 | 2.29 ± 0.22 | 0.561 |
| Mg, mmol/L | 0.99 (0.89 - 1.10) | 0.99 (0.96 - 1.09) | 0.346 |
| Pi, mmol/L | 2.20 (1.56 - 2.37) | 1.74 (1.54 - 2.14) | 0.765 |
| GLU, mmol/L | 5.48 (4.67 - 6.68) | 6.00 (4.85 - 9.68) | 0.336 |
| BUN, mmol/L | 22.22 (19.21 - 31.26) | 24.52 (14.91 - 32.64) | 0.570 |
| CREA, µmol/L | 953.33 ± 342.47 | 850.30 ± 376.79 | 0.659 |
| CysC, mg/L | 7.21 (6.46 - 7.65) | 6.87 (5.85 - 7.70) | 0.414 |
| eGFR, mL/min·per 1.73m$^2$ | 4.92 (3.66 - 6.73) | 5.05 (4.44 - 7.59) | 0.772 |
| PALB, mg/dL | 310.44 (288.15 - 331.88) | 293.39 (237.12 - 375.96) | 0.126 |
| TP, g/L | 65.49 ± 3.75 | 66.71 ± 6.14 | 0.623 |
| ALB, g/L | 37.89 ± 2.27 | 38.49 ± 3.92 | 0.605 |
| GLOB, g/L | 27.64 ± 4.06 | 28.18 ± 4.01 | 0.835 |
| A/G | 1.41 ± 0.26 | 1.39 ± 0.22 | 0.892 |
| TBIL, µmol/L | 9.61 (7.11 - 11.18) | 9.29 (8.15 - 11.53) | 0.556 |
| DBIL, µmol/L | 1.62 (1.20–2.28) | 1.72 (1.42–2.11) | 0.983 |
| ALT, U/L | 12.56 (9.70 - 16.00) | 10.29 (7.02 - 14.29) | 0.217 |
| AST, U/L | 15.14 ± 6.64 | 13.08 ± 5.15 | 0.551 |
| ALP, U/L | 86.91 (57.06 - 105.38) | 70.38 (59.16 - 99.00) | 0.715 |
| GGT, U/L | 15.90 (11.35 - 22.65) | 17.70 (10.30 - 24.40) | 0.442 |
| ApoA, g/L | 118.42 ± 23.52 | 112.52 ± 20.71 | 0.232 |
| ApoB, g/L | 65.49 (58.75 - 78.35) | 69.49 (48.12 - 91.43) | 0.729 |
| CHOL, mmol/L | 3.74 ± 0.83 | 3.75 ± 1.03 | 0.658 |
| TG, mmol/L | 1.31 (1.01 - 1.93) | 1.20 (0.73 - 2.02) | 0.676 |
| HDLC, mmol/L | 1.04 ± 0.28 | 1.02 ± 0.22 | 0.417 |
| LDLC, mmol/L | 2.28 ± 0.59 | 2.23 ± 0.81 | 0.441 |
| *SNRK* | 0.92 (0.65 - 1.21) | 0.89 (0.49 - 1.04) | 0.024 |
| *SNRK-AS1* | 1.05 (0.76 - 1.21) | 1.27 (0.84 - 1.71) | <0.001 |

Values are expressed as mean ± SD or median (25th to 75th quartiles) for continuous variables and n (%) for categorical variables, respectively. Statistical significance was assessed using Student's t test or nonparametric Mann Whitney U test. Abbreviations are defined as in Supplemental Table 1.

**Supplemental Table 5.** Multiple linear regression analysis of association of *SNRK-AS1* and clinical factors and thoracic artery calcification Agatston Score adjusted for age and sex in external validation cohort

| **Variable** | **Coefficient (B)** | **95% CI** | | | ***p* value** |
|---|---|---|---|---|---|
| Gender | 13.732 | -5.094 | to | 22.370 | **0.002** |
| Age, years | 1.677 | 1.326 | to | 2.028 | **<0.001** |
| SBP, mmHg | 0.238 | 0.064 | to | 0.412 | **0.008** |
| Pi, mmol/L | 4.643 | -2.758 | to | 12.044 | 0.219 |
| GLU, mmol/L | 0.347 | -0.760 | to | 1.454 | 0.540 |
| eGFR, mL/min·per 1.73m$^2$ | -0.328 | -0.585 | to | -0.071 | **0.013** |
| ALB, g/L | 1.972 | 0.882 | to | 3.062 | **<0.001** |
| ALT, U/L | -0.360 | -0.730 | to | 0.010 | 0.058 |
| ALP, U/L | -0.008 | -0.108 | to | 0.092 | 0.874 |
| TG, mmol/L | 7.109 | 1.484 | to | 12.734 | **0.014** |
| HDLC, mmol/L | -9.618 | -27.770 | to | 8.534 | 0.299 |
| LDLC, mmol/L | 8.498 | 2.434 | to | 14.562 | **0.006** |
| *SNRK-AS1* | 97.664 | 89.246 | to | 106.082 | **<0.001** |

Abbreviations are defined as in Supplemental Table 1, the value of thoracic artery calcification Agatston scores was taken as a square root transformation for analysis, and the regression residuals are normally distributed.

**Supplemental Table 6.** Multiple linear regression analysis of association of *SNRK* and clinical factors and thoracic artery calcification Agatston Score adjusted for age and sex in external validation cohort

| **Variable** | **Coefficient (B)** | **95% CI** | | | ***p* value** |
|---|---|---|---|---|---|
| Gender | 18.091 | 7.290 | to | 28.892 | **0.001** |
| Age, years | 1.696 | 1.271 | to | 2.121 | **<0.001** |
| SBP, mmHg | 0.128 | -0.091 | to | 0.346 | 0.251 |
| Pi, mmol/L | 7.045 | -2.152 | to | 16.242 | 0.133 |
| GLU, mmol/L | 0.011 | -1.368 | to | 1.390 | 0.988 |
| eGFR, mL/min·per 1.73m$^2$ | -0.238 | -0.564 | to | 0.088 | 0.152 |
| ALB, g/L | 0.134 | -1.209 | to | 1.478 | 0.845 |
| ALT, U/L | -0.701 | -1.166 | to | -0.235 | **0.003** |
| ALP, U/L | -0.075 | -0.199 | to | 0.049 | 0.238 |
| TG, mmol/L | 4.439 | -2.616 | to | 11.493 | 0.217 |
| HDLC, mmol/L | -2.774 | -25.135 | to | 19.588 | 0.808 |
| LDLC, mmol/L | 15.227 | 7.720 | to | 22.735 | **<0.001** |
| *SNRK* | -63.777 | -75.953 | to | -51.601 | **<0.001** |

Abbreviations are defined as in Supplemental Table 1, the value of thoracic artery calcification Agatston scores were taken as a square root transformation for analysis, and the regression residuals are normally distributed.

**Supplemental Table 7.** List of primer sequences used for Real-Time PCR and siRNA

| | **Sequences (5' – 3')** |
|---|---|
| **Plasmid** | |
| *SNRK*-SalI-F | ACGTCGACCATGGCAGGATTTAAGCGAGGG |
| *SNRK*-NotI-R | TGGCGGCCGCGATGACATGGCAACAGCTGGC |
| *SNRK*-1-269-F | ACGTCGACCATGGCAGGATTTAAGCGAGGG |
| *SNRK*-1-269-R | AAGCGGCCGCAAGCCAAGGATGATTTTCAATCTC |
| *SNRK*-1-350-F | ACGTCGACCATGGCAGGATTTAAGCGAGGG |
| *SNRK*-1-350-R | AAGCGGCCGCTGCAGATCTGGTCTGTATTTCTTTC |
| *SNRK*-270-765-F | ACGTCGACCATGCAGGGAGTGGACCCTTCAC |
| *SNRK*-270-765-R | AAGCGGCCGCGATGACATGGCAACAGCTGG |
| *SNRK*-1-430-F | ACGTCGACCATGGCAGGATTTAAGCGAGGG |
| *SNRK*-1-430-R | AAGCGGCCGCTGCGGGTGGCACCGTAGA |
| *SNRK*-351-765-F | ACGTCGACCATGAGCCCGAGCAATATCAAGG |
| *SNRK*-351-765-R | AAGCGGCCGCGATGACATGGCAACAGCTGG |
| *Snrk*-SalI-F | TCGTCGACCATGGCAGGGTTCAAGCGA |
| *Snrk*-NotI-R | TTGCGGCCGCGATGACATGGCAGCAGCTGGCGGGG |
| **Real-Time PCR** | |
| *SNRK-AS1*-F | CATCACAATGAACTTACTGGGACA |
| *SNRK-AS1*-R | GTGGTGTTCAGGTTTTGCTTTG |
| *SNRK*-F | ATGGCAGGATTTAAGCGAGGG |
| *SNRK*-R | GCCTGGCAAGTTTAACCACG |
| *RUNX2*-F | CTTCACAAATCCTCCCCAG |
| *RUNX2*-R | GAATGCGCCCTAAATCA |
| *MSX2*-F | CGGAAAATTCAGAAGATGGAGCG |
| *MSX2*-R | CGGCTTCCGATTGGTCTTGTGT |
| *SOX9*-F | AGGAAGCTCGCGGACCAGTAC |
| *SOX9*-R | GGTGGTCCTTCTTGTGCTGCAC |
| *ALPL*-F | GCTGTAAGGACATCGCCTACCA |
| *ALPL*-R | CCTGGCTTTCTCGTCACTCTCA |
| *SPP1*-F | CGAGGTGATAGTGTGGTTTATGG |
| *SPP1*-R | GCACCATTCAACTCCTCGCTTTC |
| *COL1A1*-F | GATTCCCTGGACCTAAAGGTGC |
| *COL1A1*-R | AGCCTCTCCATCTTTGCCAGCA |
| *CTNNB1*-F | CACAAGCAGAGTGCTGAAGGTG |
| *CTNNB1*-R | GATTCCTGAGAGTCCAAAGACAG |
| *GAPDH*-F | GGAAGCTTGTCATCAATGGAAATC |
| *GAPDH*-R | TGATGACCCTTTTGGCTCCC |
| **siRNA** | |
| *SNRK-AS1*-F1 | GCAUAUAGAUACUAGACUATT |
| *SNRK-AS1*-R1 | UAGUCUAGUAUCUAUAUGCTT |
| *SNRK-AS1*-F2 | GGAAUAGGUUAAAGCAAGATT |
| *SNRK-AS1*-R2 | UCUUGCUUUAACCUAUUCCTT |
| *SNRK-AS1*-F3 | GGAAGUUCAUAAGGAAGAATT |
| *SNRK-AS1*-R3 | UUCUUCCUUAUGAACUUCCTT |

### Supplemental Figures and Figure Legends

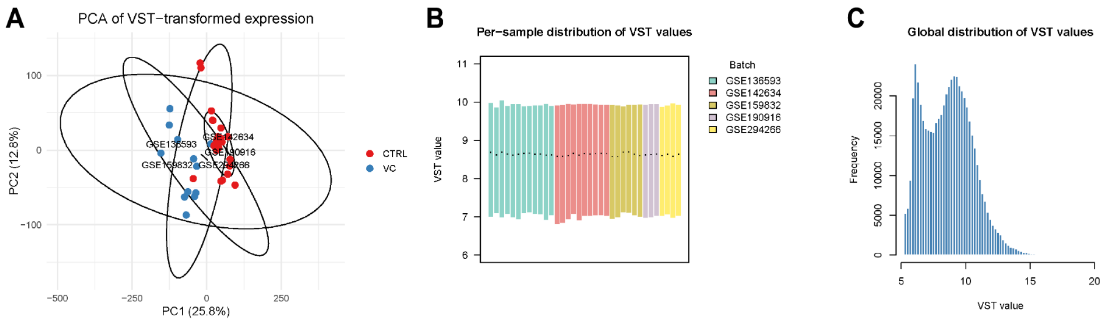


**Supplemental Figure 1. Global assessment of variance-stabilized RNA-seq expression across integrated datasets** (**A**) PCA of VST-transformed expression in mouse aorta samples, colored by group and annotated by GEO batches. (**B**) Per-sample boxplots of VST values, with samples colored by batch to visualize between-batch distribution differences. (**C**) Global histogram of all VST-transformed values across genes and samples.

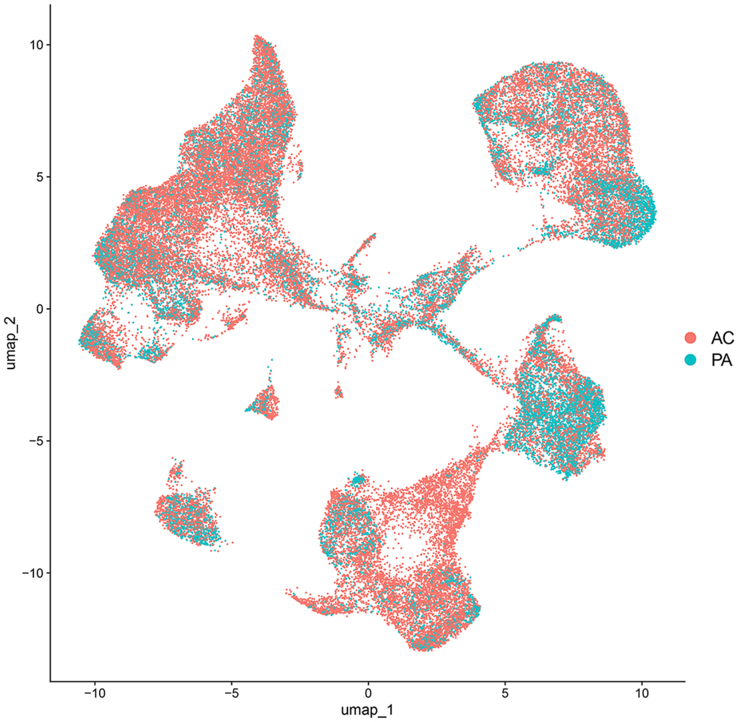


**Supplemental Figure 2. UMAP visualization of AC and PA cells after batch effect correction.**

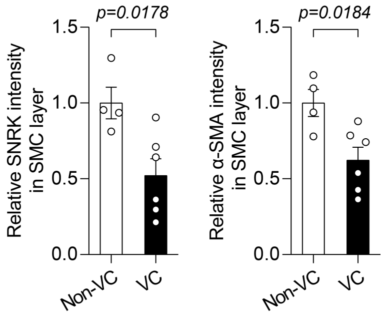


**Supplemental Figure 3. Decreased SNRK and contractile VSMC marker α-SMA level in calcified human femoral arteries.** Quantification of immunofluorescence staining of SNRK and α-SMA in non-calcified (Non-VC) or calcified (VC) human femoral artery (n=4-6). Results are presented as mean ± SEM and analyzed using two-tailed unpaired Student's *t*-test.

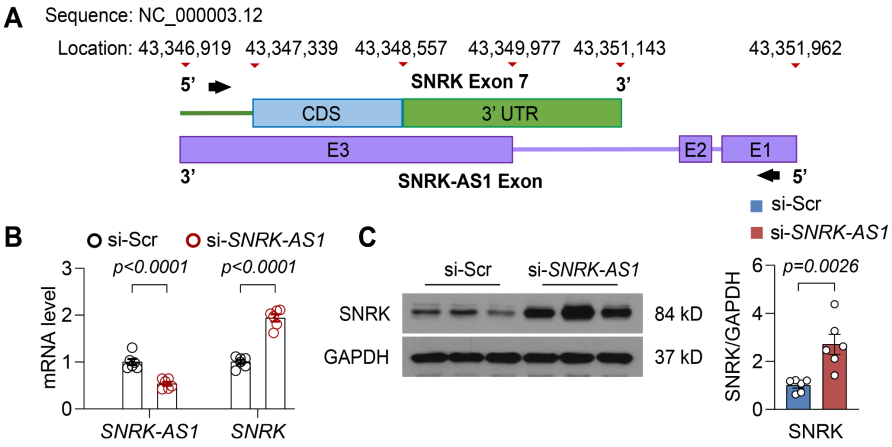


**Supplemental Figure 4. SNRK is a target for *SNRK-AS1*.** (**A**) Schematic illustration of the *SNRK-AS1*/*SNRK* Exon 7 locus. The CDS region in the SNRK Exon 7 was marked as blue. 3' UTR region was marked as green. The 3 exons for *SNRK-AS1* were marked as purple. (**B**) The expression of genes in HASMC transfected *SNRK-AS1* siRNA. (**C**) Western blot analysis of SNRK expression in HASMC transfected *SNRK-AS1* siRNA (n = 6). Results are presented as mean ± SEM. *P* values are assessed using two-way ANOVA with Sidak's post hoc test for **B** and two-tailed unpaired Student's *t*-test used for **C**.

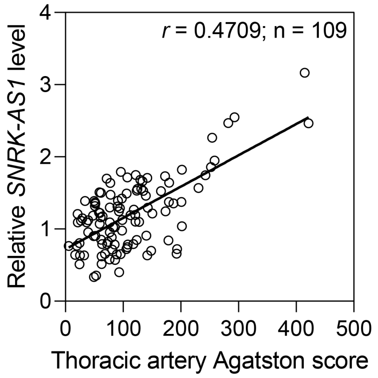


**Supplemental Figure 5**. **The correlation of PBMC *SNRK-AS1* level and square rooted thoracic artery Agatston score in CKD patients from external validation cohort.** Results are analyzed by Spearman's rank correlation coefficient test ($r$ = 0.4709, 95% CI 0.278-0.624).

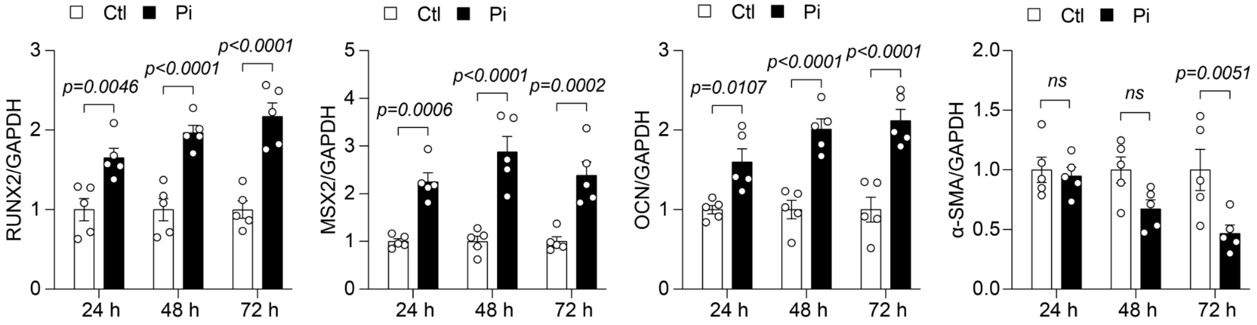


**Supplemental Figure 6. Increased RUNX2/MSX2/OCN and decreased α-SMA level in HASMC in response to calcifying stimuli.** Western blot analysis of RUNX2, MSX2, OCN, and α-SMA in human VSMC (HASMC) treated phosphate (Pi) for indicated time (n = 5). Results are presented as mean ± SEM, and analyzed using two-tailed unpaired Student's $t$-test.

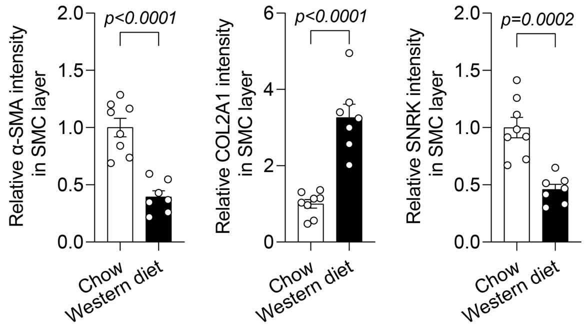


**Supplemental Figure 7. Reduced α-SMA, increased COL2A1 level, and SNRK in calcified mice arteries.** Quantification of immunofluorescence staining of α-SMA, COL2A1, and SNRK in brachiocephalic artery of *Apoe*$^{-/-}$ mice fed with chow diet or western diet for 24 weeks (n = 7-8). Results are presented as mean ± SEM and analyzed using two-tailed unpaired Student's *t*-test.

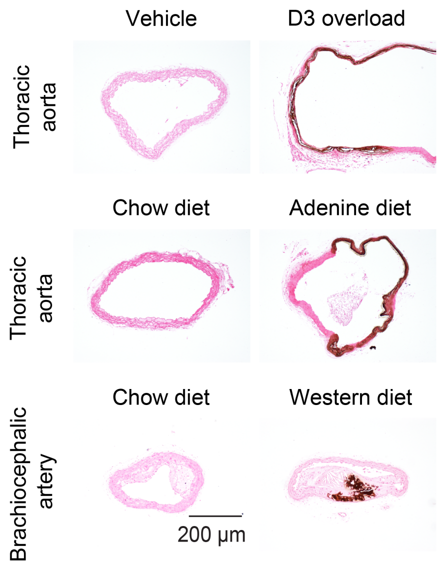


**Supplemental Figure 8. Vascular calcification induction in three mouse models.** Representative von Kossa staining of thoracic aorta and brachiocephalic artery of indicated mouse models of vascular calcification (n = 6).

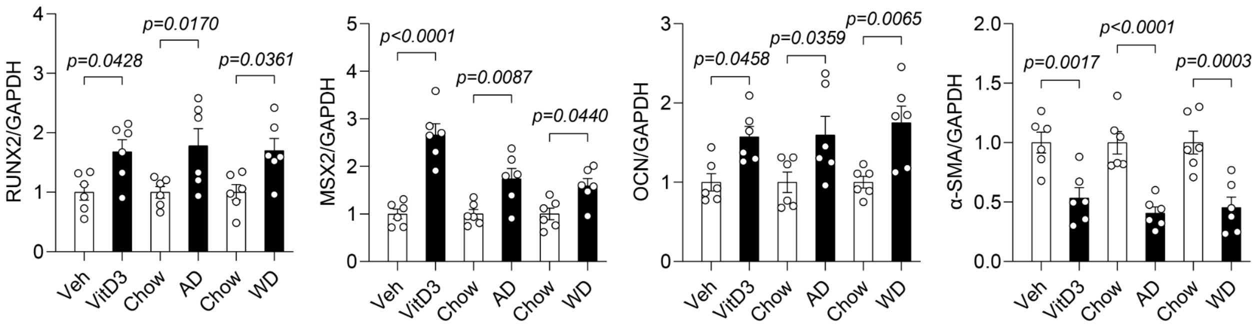


**Supplemental Figure 9. Increased RUNX2/MSX2/OCN and decreased α-SMA level in aorta of three mouse models of calcification.** Western blot analysis of RUNX2, MSX2, OCN, and α-SMA in aortic SMC layer of indicated mice (n = 6). Vitamin D3 (D3), Adenine diet (AD), Western diet (WD). Results are presented as mean ± SEM and analyzed using two-tailed unpaired Student's *t*-test.

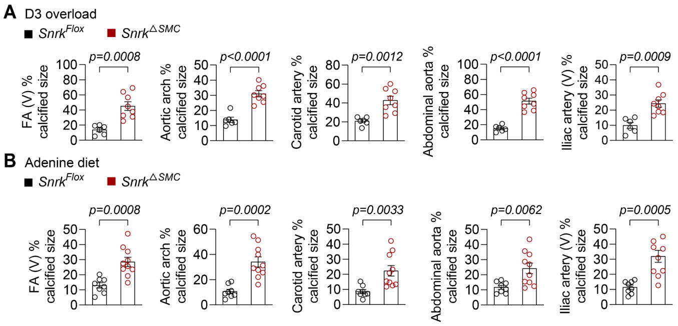


**Supplemental Figure 10. VSMC-specific SNRK deficiency promotes vascular calcification.** Calcification size analysis of femoral artery (V), aortic arch, carotid artery, abdominal aorta, and iliac artery (V) of vitamin D3 overload-induced (**A**) and adenine diet-induced (**B**) mouse models of vascular calcification (n = 6-10). Vertical section (V). Results are presented as mean ± SEM, and analyzed using unpaired *t*-test with Welch's correction.

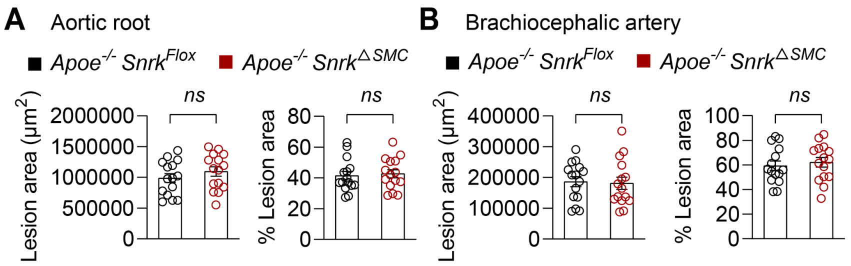


**Supplemental Figure 11. SNRK deficiency has no effect on plaque formation in situ.** (**A** and **B**) Quantification of atherosclerotic lesion size and percentage of lesion size within lumen area in aortic root and brachiocephalic artery of *Apoe*$^{-/-}$ *Snrk*$^{Flox}$ or *Apoe*$^{-/-}$ *Snrk*$^{\triangle SMC}$ mice fed with western diet (n = 15). Results are presented as mean ± SEM and analyzed using two-tailed unpaired Student's *t*-test.

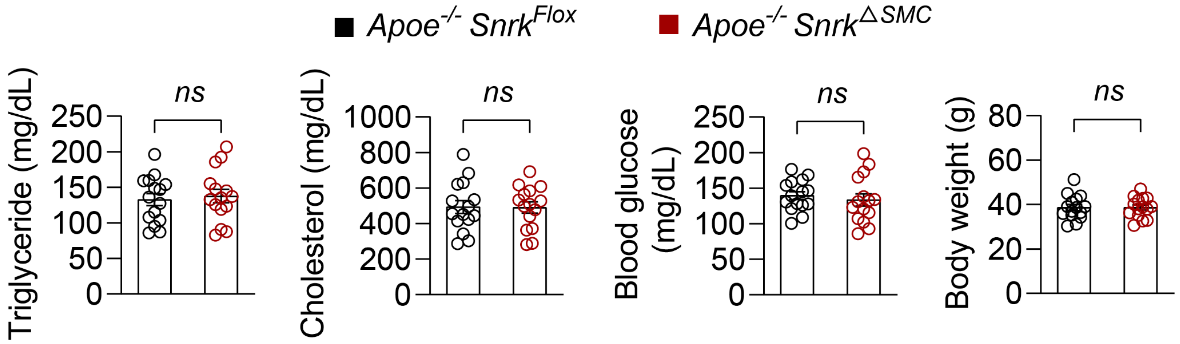


**Supplemental Figure 12. SNRK deficiency has no effect on metabolic parameters.** Biochemical measurement of serum triglyceride, cholesterol, blood glucose, and body weight of *Apoe*$^{-/-}$ *Snrk*$^{Flox}$ or *Apoe*$^{-/-}$ *Snrk*$^{\triangle SMC}$ mice fed with western diet (n = 15). Results are presented as mean ± SEM and analyzed using two-tailed unpaired Student's *t*-test.

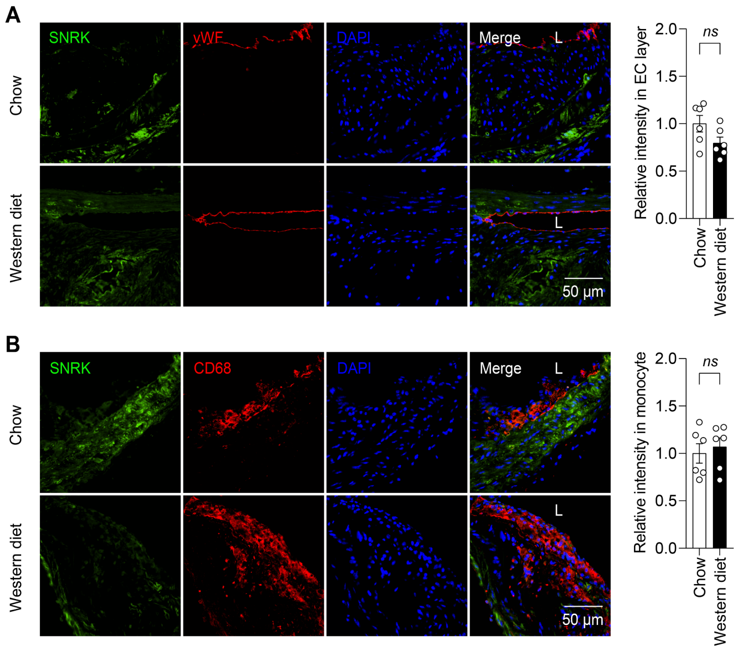


**Supplemental Figure 13. SNRK level in endothelial and macrophage of calcified mice arteries.** Quantification and immunofluorescence staining of SNRK co-located with vWF (**A**) or CD68 (**B**) in brachiocephalic artery of *Apoe*$^{-/-}$ mice fed with chow diet or western diet for 24 weeks (n = 6). Results are presented as mean ± SEM and analyzed using two-tailed unpaired Student's *t*-test.

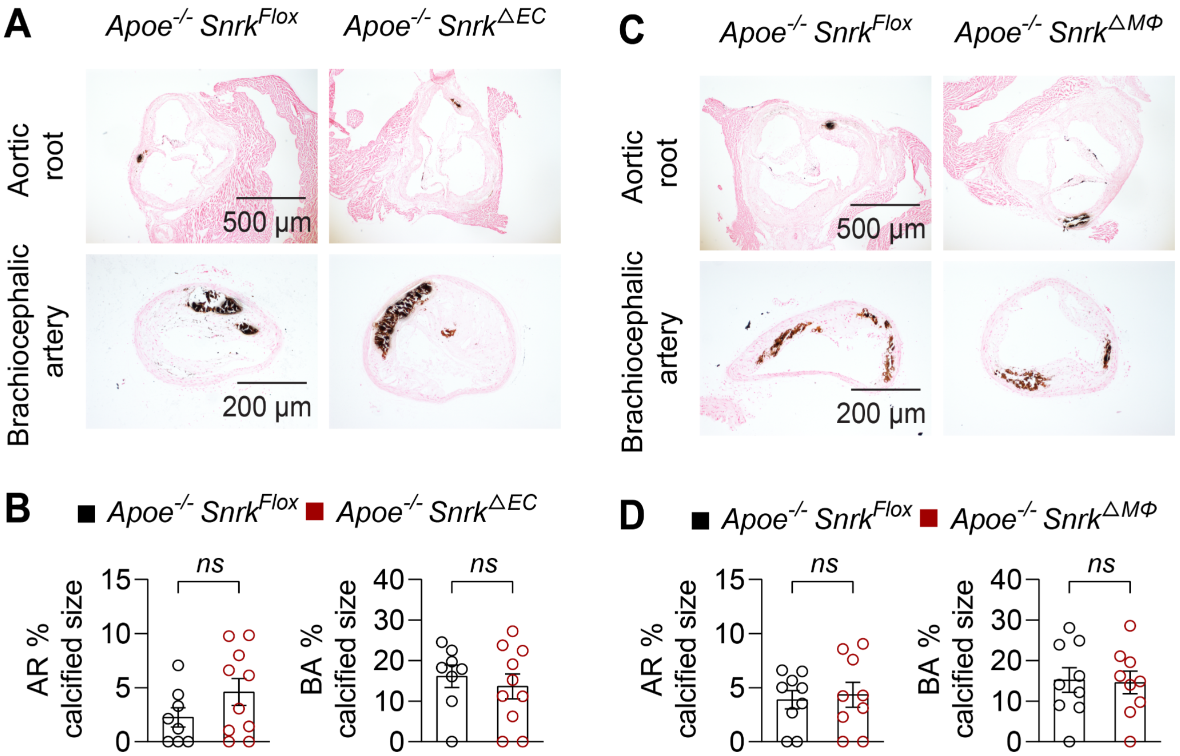


**Supplemental Figure 14. SNRK deficiency in endothelial or macrophage has mild effect on calcification formation.** Von Kossa staining and calcification size analysis of aortic root (AR) and brachiocephalic artery (BA) of *Apoe*$^{-/-}$ *Snrk*$^{Flox}$ or endothelial-specific SNRK deletion (*Apoe*$^{-/-}$ *Snrk*$^{\triangle EC}$) mice (**A** and **B**) , and *Apoe*$^{-/-}$ *Snrk*$^{Flox}$ or macrophage-specific SNRK deletion (*Apoe*$^{-/-}$ *Snrk*$^{\triangle M\Phi}$) mice (**C** and **D**) fed with western diet for 24 weeks (n = 8-10). Results are presented as mean ± SEM and analyzed using two-tailed unpaired Student's *t*-test.

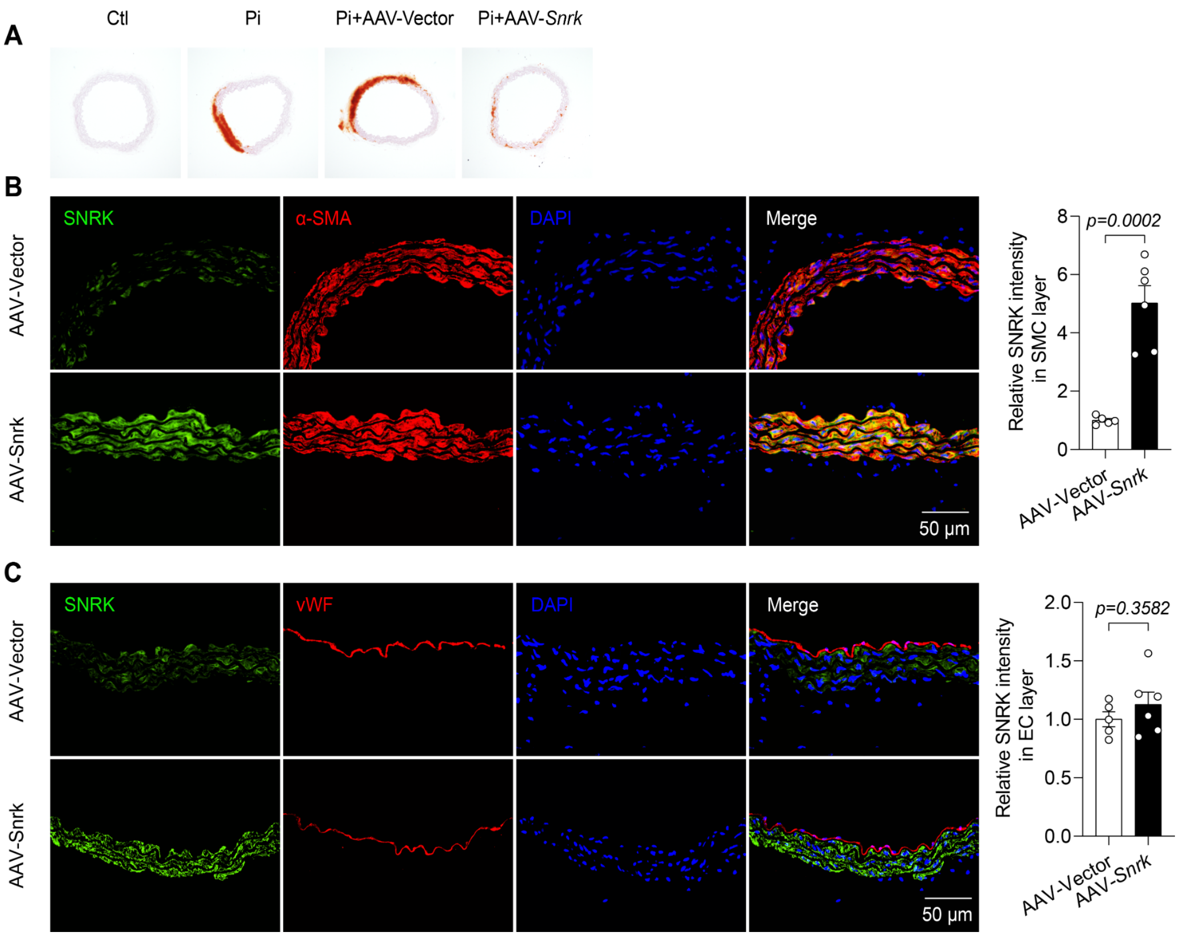


**Supplemental Figure 15. SNRK overexpression in aortic rings.** (**A**) Alizarin red S staining of aortic rings isolated from C57BL/6J mice infected vector or AAV-*Snrk* (n = 6). (**B** and **C**) Immunofluorescence staining of SNRK co-located with α-SMA or vWF in aorta isolated from C57BL/6J mice infected vector or AAV-*Snrk* (n = 5-6). Results are presented as mean ± SEM and analyzed using two-tailed unpaired Student's *t*-test.

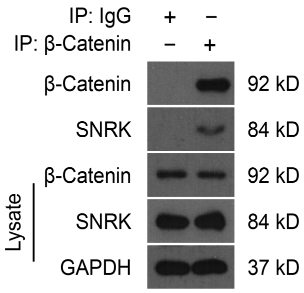


**Supplemental Figure 16. SNRK associates with endogenous β-Catenin.** Co-IP assay of endogenous SNRK and β-Catenin in HASMCs (n = 3).

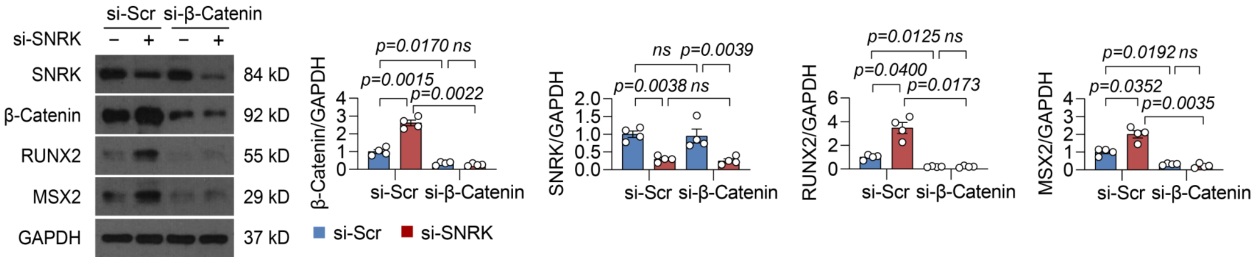


**Supplemental Figure 17. β-Catenin silencing eliminated the upregulation of RUNX2 and MSX2 levels induced by SNRK deficiency.** Western blot analysis of β-Catenin, SNRK, RUNX2, and MSX2 in HASMCs transfected SNRK siRNA or/and β-Catenin siRNA for 24 hours (n = 4). Results are presented as mean ± SEM. *P* values are assessed using Welch's ANOVA with Dunnett's post hoc test.

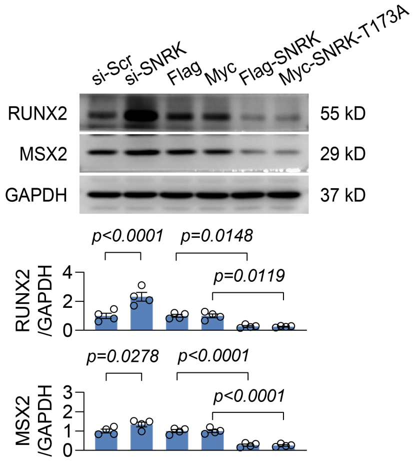


**Supplemental Figure 18. Calcification-related factors expression in HASMC with SNRK depletion or overexpression.** Western blot analysis of RUNX2 and MSX2 in HASMC transfected SNRK siRNA or SNRK plasmid as indicated (n = 4). Results are presented as mean ± SEM and analyzed using two-tailed unpaired Student's *t*-test.

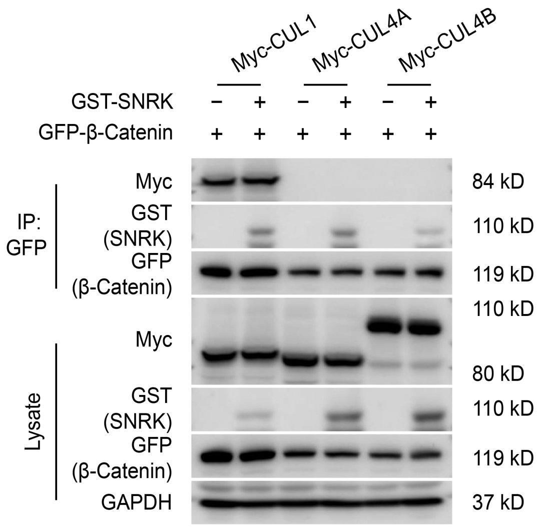


**Supplemental Figure 19. SNRK promotes β-Catenin degradation through interacting with Cul1 SCF complex.** Western blot analysis of HASMC transfected GST-SNRK, Myc-Cul1, Myc-Cul4A or Myc-Cul4B, and GFP-β-Catenin plasmid as indicated (n=4).

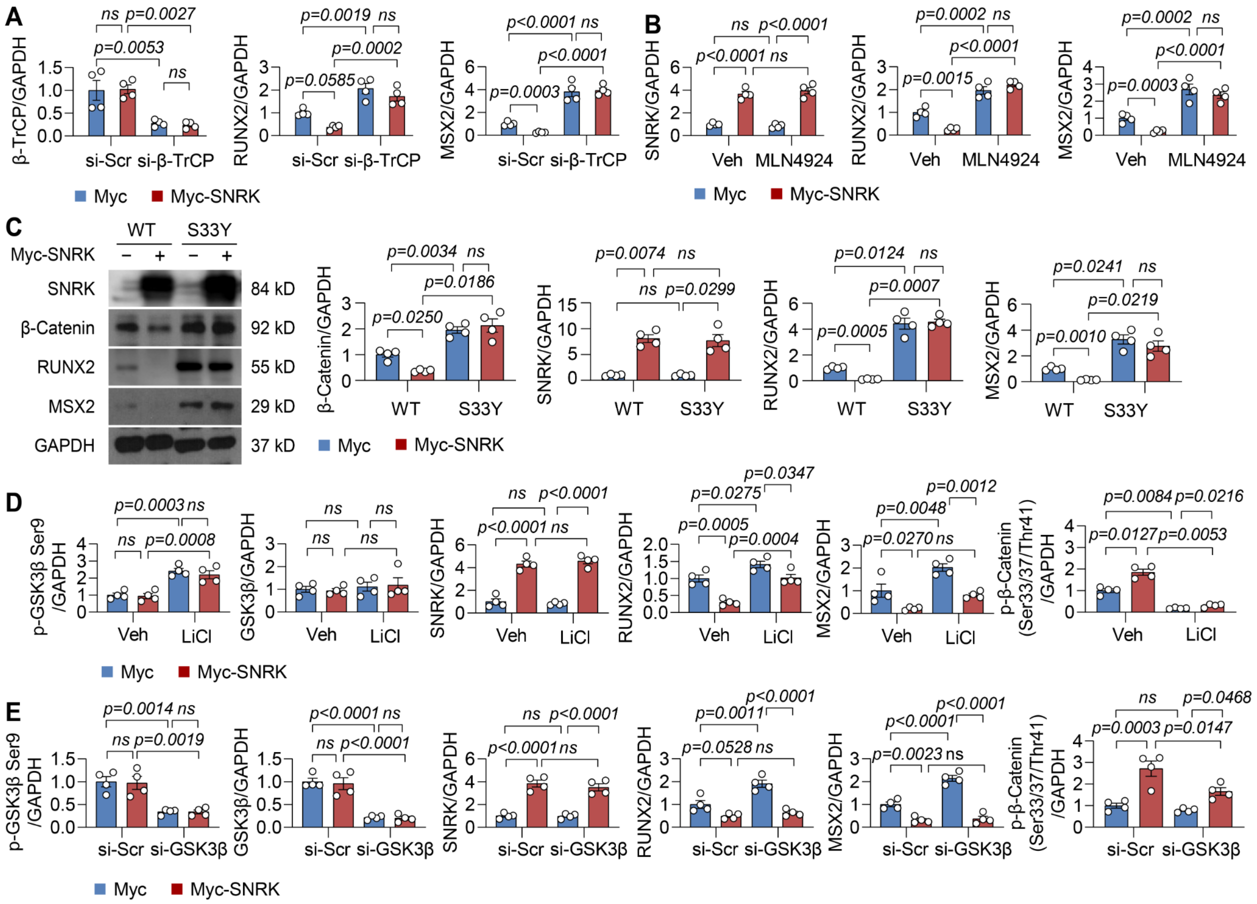


**Supplemental Figure 20. SNRK directly targets β-Catenin degradation in a destruction complex-independent manner.** (**A**) Western blot analysis of β-TrCP, RUNX2, and MSX2 in HASMC transfected with β-TrCP siRNA or SNRK plasmid as indicated (n = 4). (**B**) Western blot analysis of SNRK, RUNX2, and MSX2 in HASMC transfected SNRK and treated with vehicle or SCF complex inhibitor MLN4924 (1 μmol/L) for 6 hours (n = 4). (**C**). Western blot analysis of β-Catenin, SNRK, RUNX2, and MSX2 in HASMCs transfected SNRK and wild type

(WT) β-Catenin or β-Catenin with S33Y mutation for 48 hours (n = 4). (**D**) Western blot analysis of p-GSK3β Ser9, GSK3β, SNRK, RUNX2, and MSX2 in HASMC transfected SNRK and treated with vehicle or GSK3β inhibitor LiCl (20 mmol/L) for 24 hours (n = 4). (**E**) Western blot analysis of p-GSK3β Ser9, GSK3β, SNRK, RUNX2, and MSX2 in HASMC transfected GSK3β siRNA or SNRK plasmid as indicated (n = 4). Results are presented as mean ± SEM. *P* values are assessed using one-way ANOVA with Sidak's post hoc test.

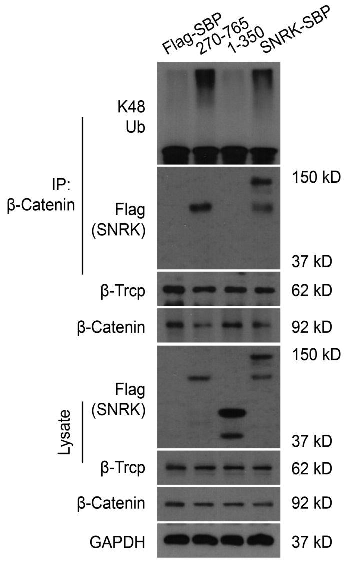


**Supplemental Figure 21. The aa 270-765 SNRK domain bridges endogenous β-Catenin and β-TrCP.** Co-IP assay of endogenous β-Catenin in HASMCs transfected full length SNRK plasmid or truncated SNRK plasmid (n = 3).

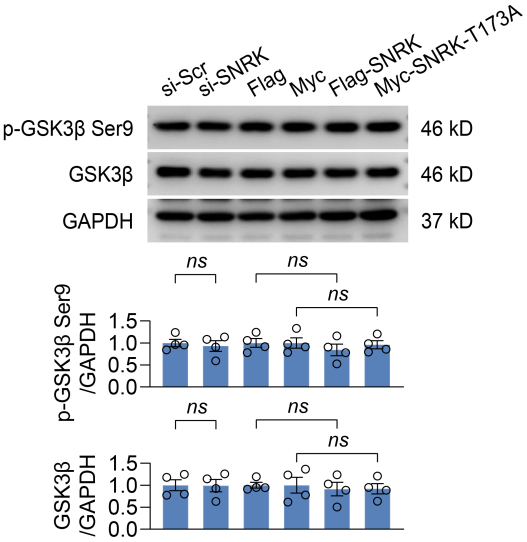


**Supplemental Figure 22. Phosphorated GSK3β levels in HASMC with SNRK depletion or overexpression.** Western blot analysis of p-GSK3β Ser9 and GSK3β in HASMC transfected SNRK siRNA or SNRK plasmid as indicated (n = 4). Results are presented as mean ± SEM and analyzed using two-tailed unpaired Student's *t*-test.

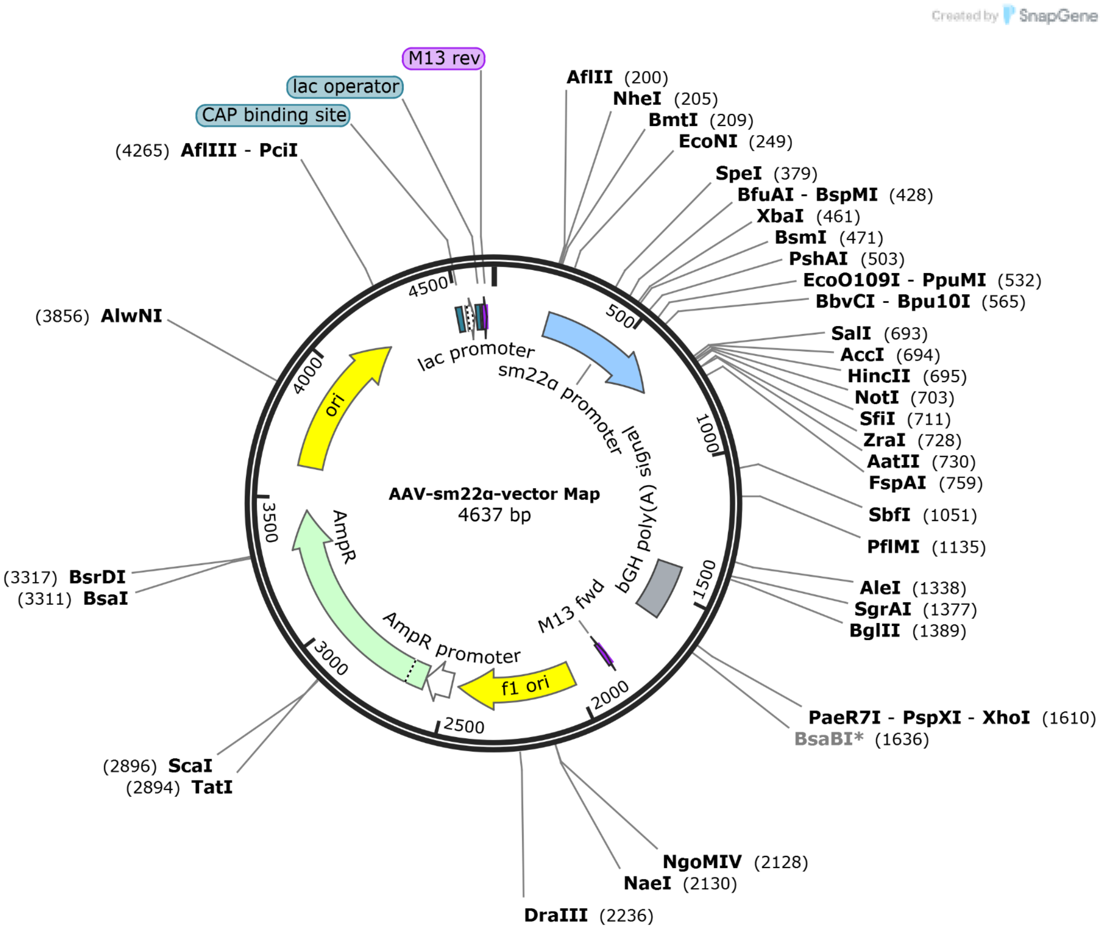


**Supplemental Figure 23. Annotated vector map of the AAV construct containing the *Tagln* (SM22α) promoter.**

8. Wu S, Lu Q, Wang Q, Ding Y, Ma Z, Mao X, Huang K, Xie Z, Zou MH. Binding of FUN14 Domain Containing 1 With Inositol 1,4,5-Trisphosphate Receptor in Mitochondria-Associated Endoplasmic Reticulum Membranes Maintains Mitochondrial Dynamics and

Function in Hearts in Vivo. *Circulation*. 2017;136:2248-2266. doi: 10.1161/CIRCULATIONAHA.117.030235

1289. doi: 10.1038/s41591-019-0512-5

15. Alsaigh T, Evans D, Frankel D, Torkamani A. Decoding the transcriptome of calcified atherosclerotic plaque at single-cell resolution. *Commun Biol*. 2022;5:1084. doi: 10.1038/s42003-022-04056-7
